# Peptide-induced hydration of lipid bilayers modulates packing pattern and conformations of hydrocarbon chains – a potential pathway for peptide translocation?

**DOI:** 10.1101/2025.02.04.636480

**Authors:** Lea Pašalić, Barbara Pem, Andreja Jakas, Ana Čikoš, Nikolina Groznica, Tihana Mlinarić, Matilde Accorsi, Agustín Mangiarotti, Rumiana Dimova, Danijela Bakarić

**Author notes:** Corresponding author.; (Danijela Bakarić).

## Abstract

Cell-penetrating peptides (CPPs) with a cationic-hydrophobic character are recognized as carriers for delivering various therapeutics and diagnostic agents across cell membranes and into the cells. Among the most studied CPPs, nona-arginine (R9) exhibits superior penetration compared to nona-lysine (K9), suggesting that the penetration ability depends not only on charge, distribution and concentration of peptides but also on the lipid membrane composition. However, for heptapeptides composed of arginine (R), lysine (K) and phenylalanine (F) residues, which show some CPPs properties, these interactions remain unexplored. This study sheds light on the adsorption of R5F2/K5F2 on model prokaryotic (PRO) and eukaryotic (EU) lipid membranes containing a zwitterionic lipid (phosphatidylcholine; PC) and an anionic lipid (either phosphatidylglycerol, PG, in the PRO model, or phosphatidylserine, PS in EU) at the 90:10 molar ratio. Using differential scanning calorimetry (DSC) and temperature-dependent UV-Vis spectroscopy, we observed peptide-induced changes in the interfacial water layer that affect the fluidity and rigidity of lipid bilayers. The distinct adsorption behavior of R5F2/K5F2 on PRO and EU lipid bilayers revealed the changes in lipid packing and hydrocarbon chain conformations as exclusively peptide-dependent features. The peptide-induced formation of vacancies in the non-polar bilayer part is consistent with partial leakage observed in giant unilamellar vesicles. The synchronized arrangement could represent a mechanism for the concerted translocation of CPPs, along with their potential cargo across the lipid membrane. This study provides new insights into the peptide-lipid interactions underlying CPPs functionality.

## 1. Introduction

As one of the most fundamental parts of the cell, the plasma membrane serves as an efficient barrier between the cell interior and its environment, regulating the movement of substances in and out of the cell [1,2]. However, this protective function also prevents therapeutic agents from easily penetrating the membrane, challenging drug delivery [3]. Therefore, a significant effort has been invested to develop delivery carriers capable of efficiently crossing membranes and transporting biologically active cargoes into cells. Recently, attention has shifted to cell-penetrating peptides (CPPs), which consist of no more than 30 amino acids and are characterized by a high proportion of positively charged amino acids, such as arginine (Arg/R) and lysine (Lys/K) [4,5]. These peptides have the remarkable ability to cross cell membranes and deliver various types of cargo [6,7]. The first discovered CPP was derived from Transactivating Transcriptional Activator (TAT), a protein of the human immunodeficiency virus 1 (HIV-1), identified in the late 1980s [8,9]. Initially, it was found that the full-length protein crossed the plasma membrane, while later, small fragments able to penetrate the membrane were identified [7]. More specifically, it was found that the R-and K-rich sequence consisting of eleven amino acids (GRKKRRQRRRC, where G is glycine, Q is glutamine and C is cysteine) is more efficient than the full-length protein [10].

Several studies demonstrated that CPPs with R residues are far more efficient than their K counterparts, regardless of having equal net charge [11–13]. More specifically, Robison *et al*. demonstrated that R9 interacts more strongly than K9 with bilayers composed dominantly of 1-palmitoyl-2-oleoyl-*sn*-glycero-3-phosphocholine (POPC) and containing a variable amount of 1-palmitoyl-2-oleoyl-*sn*-glycero-3-phospho-(1’-rac-glycerol) (sodium salt) (POPG) [12]. This effect can be attributed to the distinct chemical properties of the cationic groups in R and K; in particular, the guanidinium group (Gdm^+^) in R forms a bidentate interaction with the phosphate on the lipid headgroup, while the amino group (NH_3_^+^) of K can only interact with a single headgroup at a time [14–16]. Furthermore, the adsorption of R9 and K9 seems to be strongly influenced by factors such as peptide concentration [14,17], ionic strength [18,19], the composition of the surrounding aqueous environment [19], and the lipid membrane composition [20,21]. Interestingly, Kamat *et al.* demonstrated that even shorter peptides of mixed cationic hydrophobic character adsorbed at the bilayer surface may display CPP features. Having compartmentalized negatively charged RNA molecules in large unilamellar liposomes (LUVs) constituted from POPC lipids, one of the crucial findings that emerged from conducted research is that some R-and phenylalanine (Phe/F)-based heptapeptides are capable of traversing the lipid membrane [22]. To elucidate the adsorption fashion of R5F2 and K5F2 peptides on the surface of lipid bilayers composed from dominantly 1,2-dipalmitoyl-*sn*-glycero-3-phosphocholine (DPPC), as well as to identify the features that make R-based CPPs more efficient than K-based analogs, recently we observed increased membrane interactions with R5F2 compared to K5F2, and predicted the stronger attachment of the former to the polar groups of the lipid [23]. The easier deprotonation of NH_3_^+^ moiety (p*K*_a_ ≈ 9.3) compared to that of Gdm^+^ moiety (p*K*_a_ ≈ 13.6) as constitutive elements of K and R, respectively [15], appears to be rather important in the dependence of the internalization of (potential) CPPs on lipid membrane composition [24–26]. For instance, despite bearing a negative net charge, the composition of plasma membranes significantly varies across prokaryotic and eukaryotic organisms [2,27], resulting in distinct physicochemical properties [28].

To gain a deeper insight into how certain peptides adsorb at the surface of membranes having different charge distributions, in this paper we explored the interaction pattern of R5F2 and K5F2 on lipid membranes that serve as models of prokaryotic (PRO) and eukaryotic (EU) lipid membranes represented by mixtures of DPPC and 1,2-dipalmitoyl-*sn*-glycero-3-phospho-(1′-rac-glycerol) sodium salt (DPPG) (90%:10%) and DPPC and 1,2-dipalmitoyl-*sn*-glycero-3-phosphoserine (DPPS) (90%:10%), respectively. By examining the lipids that undergo a phase change upon heating in the experimentally accessible temperature range, we highlighted the structural features inherent to the lipid bilayer arrangement in a particular phase and discerned them from those exclusively peptide-dependent. Accordingly, we employed DSC and temperature-dependent UV-Vis spectroscopy in the characterization of thermotropic properties of lipid bilayers in terms of their pretransition (*T*_p_) and the main phase transition temperatures (*T*_m_), the former being associated with the phase transition of DPPC lipids from the gel (L_β’_) to the ripple phase (P_β’_), whereas the latter with the ripple (P_β’_) to the fluid phase (L_α_) [29]. A molecular-level picture of these events is elaborated in terms of utilizing FTIR spectroscopy, giant vesicle observations and MD simulations of PRO and EU lipid bilayers, which enabled us to identify the most sensitive lipid bilayer regions on peptide adsorption, and how are they entangled with the changes in the interfacial water layer.

## 2. Experimental

### 2.1 Chemicals and preparation of large and giant unilamellar liposomes (LUVs and GUVs) in the absence/presence of peptides R5F2 and K5F2

Peptides R5F2 and K5F2 were synthesized following the solid-phase peptide synthesis protocol [23,30] (the details on their characterization are presented in Supporting Information, section S1).

*LUVs preparation.* 1,2-dipalmitoyl-*sn*-glycero-3-phosphocholine (DPPC), 1,2-dipalmitoyl-*sn*-glycero-3-phospho-L-serine sodium salt (DPPS) and 1,2-dipalmitoyl-*sn*-glycero-3-phospho-(1′-rac-glycerol) sodium salt (DPPG) were purchased as white powders from Avanti Polar Lipids (≥ 99%). The aqueous solution of phosphate buffer (PB) of ionic strength (*I* = 100 mmol dm^−3^) was prepared from commercially available sodium hydrogen phosphate, anhydrous (Na_2_HPO_4_, ≥ 99%, Kemika, Zagreb, Croatia) and sodium phosphate monobasic (NaH_2_PO_4_, p.a., Kemika, Zagreb, Croatia) in Milli-Q water and titrated with freshly prepared sodium hydroxide solution (NaOH, T.T.T., p.a.) of ionic strength (*I*(NaOH) = 100 mmol dm^−3^) to achieve pH ≈ 7.4. Stock solutions of lipids were prepared by dissolving 100 mg of DPPC/DPPG/DPPS in 10 mL of chloroform (CHCl_3_; colorless liquid, p.a., Carlo Erba) resulting in *γ*(DPPC/DPPG/DPPS) = 10 mg mL^-1^ (*c*(DPPC/DPPG/DPPS) = 0.0135 mol dm^−3^/0.0132 mol dm^−3^/0.0134 mol dm^−3^), that was further used in the preparation of multilamellar liposomes (MLV). 2.7 mL of DPPC stock solution was mixed with 0.3 mL of stock solution of DPPG/DPPS to achieve 90:10 (mol/mol) DPPC/DPPG (PRO) or DPPC/DPPS (EU) mixtures. After pipetting the mixtures into round-bottom flasks and removing CHCl_3_ on a rotary evaporator, lipid films were additionally dried under the Ar stream. Obtained lipid films were suspended in 6 mL of prepared PB to obtain MLVs composed of DPPC/DPPG (PRO) and DPPC/DPPS (EU) lipids. The preparation of both PRO and EU MLVs took place in three steps: starting with vortexing the suspensions, followed by heating in a hot H_2_O bath (60°C), and finishing with cooling in an ice bath (∼4°C). 2 mL of MLV suspensions (PRO and EU) was pipetted into the separate flasks (to examine the peptide-free suspensions). Before adding the peptides to the lipid suspension, 10 mg of R5F2/K5F2 peptides were dissolved in 10 mL of PB (pH ≈ 7.4) to achieve their mass concentration *γ*(R5F2/K5F2) = 1 mg mL^−1^ (*c*(R5F2/K5F2) = 0.00091 mol dm^−3^/0.001 mol dm^−3^). In the suspensions of MLVs that mimic PRO and EU systems 1.48 ml of R5F2 and l.29 mL of K5F2 were added to reach 1:30 peptide-to-lipid molar ratio [23]. To obtain LUVs of appropriate composition in the absence and the presence of peptides, MLV suspensions without and with peptides were pushed at least 31 times through a 100 nm size polycarbonate membrane using an Avanti® Mini Extruder with a holder/heating block (at 55 °C) and 10 mm supporting filters. Mass concentrations of LUVs without and with peptides intended for DSC and FTIR measurements were *γ* =5 mg mL^−1^, for temperature-dependent UV-Vis measurements *γ* =1 mg mL^−1^ and for DLS and ELS measurements *γ* = 0.05 mg mL^−1^, respectively.

*GUVs preparation.* 1-palmitoyl-2-oleoyl-glycero-3-phosphocholine (POPC), 1-palmitoyl-2-oleoyl-*sn*-glycero-3-phospho-L-serine (POPS) and 1-palmitoyl-2-oleoyl-*sn*-glycero-3-phospho-(1’-rac-glycerol) (POPG) were purchased as white powders from Avanti Polar Lipids (99%). Sucrose (C_12_H_22_O_11_, ≥ 99.5%) was purchased from Sigma-Aldrich. The fluorophore, 1,1’-dioctadecyl-3,3,3’,3’-tetramethlyindocarbocyanine perchlorate (DilC_18_) was purchased form Termo Fisher Scientific. Stock solutions of lipids in CHCl_3_, at *c* = 3mM, were prepared in 2 different compositons, POPC+10 mol% POPG+0.5 mol% DilC_18_ (labeled as PRO’) and POPC+10% mol POPS+0.5 mol% DilC_18_ (labeles as EU’). The peptides R5F2 and K5F2 were dissolved in PB at concentration of 200 µM. The osmolarity of the peptide solution, measured by freezing point osmometer Osmomat 3000 (Gonotec, Berlin, Germany), was fixed to approximately 225 mOsm kg^-1^. The osmolarity of the freshly prepared sucrose and glucose solutions were adjusted to match the one of the peptide solution (225 mOsm kg^−1^). Giant unilamellar vesicles (GUVs) that serve as models of prokaryotic (PRO’) and eukariotyc (EU’) lipid membranes in the absence/presence of R5F2/K5F2 were prepared by the electroformation method [31] described below and explored by confocal fluorescence microscopy. Indium tin oxide (ITO)-coated glass slides were carefully washed in the following order, starting with Milli-Q water, ethanol (EtOH; colorless liquid, p.a., Honeywell) and CHCl_3_ and followed by drying under N_2_ stream. Then, 6 µL of a stock solution of lipid in CHCl_3_ (*c* = 3 mM) was spread on the conductive side of the ITO-coated slides. The slides were placed under vacuum for one hour to ensure evaporation of the organic solvent. The two slides with the lipid-coated sides facing each other were assembled to sandwich a rectangular Teflon spacer and the electroformation chamber was filled with 2 mL of sucrose solution. The chamber was connected to a signal generator (Agilent, USA) at the voltage amplitude of 1-1.6 V_pp_ and frequency of 10 Hz. After one hour, GUVs are harvested and transferred to an Eppendorf tube.

### 2.2 Dynamic and Electrophoretic light scattering (DLS and ELS) of LUVs: Measurements and Data Analysis

The size distribution of liposomes was determined using dynamic light scattering with a photon correlation spectrophotometer (Zetasizer Nano ZS, Malvern Instruments, Worcestershire, UK) equipped with a 532 nm (green) laser. The average hydrodynamic diameter (*d*_h_) was specified as the value at the peak maximum of the volume size distribution. The results reported correspond to the average of six measurements conducted at 25 °C. The zeta potential (*ζ*-potential) was measured by ELS and calculated from the measured electrophoretic mobility through the Henry equation using the Smoluchowski approximation. The measurements were repeated three times. The DLS and ELS data processing was performed using Zetasizer software version 7.13 (Malvern Instruments). All suspensions were measured at *γ* = 0.05 mg mL^−1^.The average hydrodynamic diameter values of PRO LUVs were in the range 115 nm ≤ *d*_h_ ≤ 125 nm (no peptide), 125 nm ≤ *d*_h_ ≤ 140 nm (+R5F2), 140 nm ≤ *d*_h_ ≤ 160 nm (+ K5F2), whereas those of EU LUVs in the range 100 nm ≤ *d*_h_ ≤ 110 nm (no peptide), 120 nm ≤ *d*_h_ ≤ 130 nm (+R5F2), 110 nm ≤ *d*_h_ ≤ 115 nm (+ K5F2), respectively. Accordingly, ζ-potential values of PRO LUVs were −6.5 ± 0.4 mV (no peptide), −4.5 ± 0.9 mV (+R5F2), −10.0 ± 0.8 mV (+ K5F2), and those of EU LUVs were −16.0 ± 0.3 mV (no peptide), −13.6 ± 0.5 mV (+R5F2), −0.2 ± 0.6 mV (+ K5F2), respectively (see Supporting Information, Fig. S1).

### 2.3. Differential Scanning Calorimetry (DSC) of LUVs: Data Acquisition and Curve Analysis

The measurements of PRO and EU systems ± R5F2/K5F2 in the form of LUVs were conducted in microcalorimeter Nano-DSC, TA Instruments (New Castle, USA) using the TA Instruments Nano Analyze software package. In addition to LUVs, MLVs in the absence/presence of peptides were additionally measured so that, based on qualitative differences of associated DSC curves, we could confidently claim that LUVs do not aggregate during the measurement (Fig. S2 in Supporting Information). All suspensions and PB solution were degassed for 10 minutes before measurements. The suspensions of LUVs (and MLVs) of PRO ± R5F2/K5F2 and EU ± R5F2/K5F2 (*γ*(lipids) = 5 mg mL^−1^) were heated at a scan rate of 1 °C min^−1^ in two repeated heating–cooling cycles in a temperature range of 20–60 °C as duplicates. A reference, PB, was examined in the temperature range 10–90 °C. Thermotropic properties of PRO ± R5F2/K5F2 and EU ± R5F2/K5F2 systems were determined from a thermal history-independent second heating run [32,33]. First, the DSC curve of PB was subtracted from a DSC curve of lipid suspensions (both obtained from the second heating run); second, a temperature range extending from 30 to 52 °C was chosen for further analysis since the dominant lipid in both systems (DPPC) in this range undergo both pretransition (*T*_p_) and the main phase transition (*T*_m_), i.e. the transitions that are distinguished by the periodic undulations at the surface of the lipid bilayer and by breaking of van der Waals interactions between hydrocarbon chains, increased hydration of polar headgroup region and consequential bilayer thinning [34–38], respectively. When possible, *T*_p_ was estimated as the deviation from the baseline of the resultant DSC curve, whereas *T*_m_ was determined from the maximum of the signal. Since some explored systems displayed two maxima during melting [39–41], we designated the melting temperature obtained from DSC as *T*_m, 1(/2)_.

### 2.4 UV-Vis spectroscopy of LUVs: Acquisition of temperature-dependent spectra and data analysis

UV-Vis spectra of PRO ± R5F2/K5F2 and EU ± R5F2/K5F2 (*γ*(PRO/EU) = 1 mg mL^−1^) were measured on a UV-Vis spectrophotometer Thermo Scientific Nanodrop 2000 (Thermo Fischer Scientific, Waltham, MA, USA) within the spectral range of 250–500 nm. The samples were pipetted into 1 mL quartz cuvettes and placed in a temperature-controlled cuvette holder. The spectra of lipid suspensions in the presence/absence of R5F2/K5F2 were recorded at least three times (different cuvettes) in a temperature range 30–52 °C. The spectra of PB ± R5F2/K5F2 were collected once in the same temperature range. Obtained UV-Vis spectra were smoothed (Savitzky-Golay; polynomial of a 3^rd^ degree and 10 points) [42] and analyzed in the spectral range 250-300 nm (Fig. 2) applying multivariate curve analysis (MCA) [43] using publicly available Matlab code [44].

**Fig. 1.**
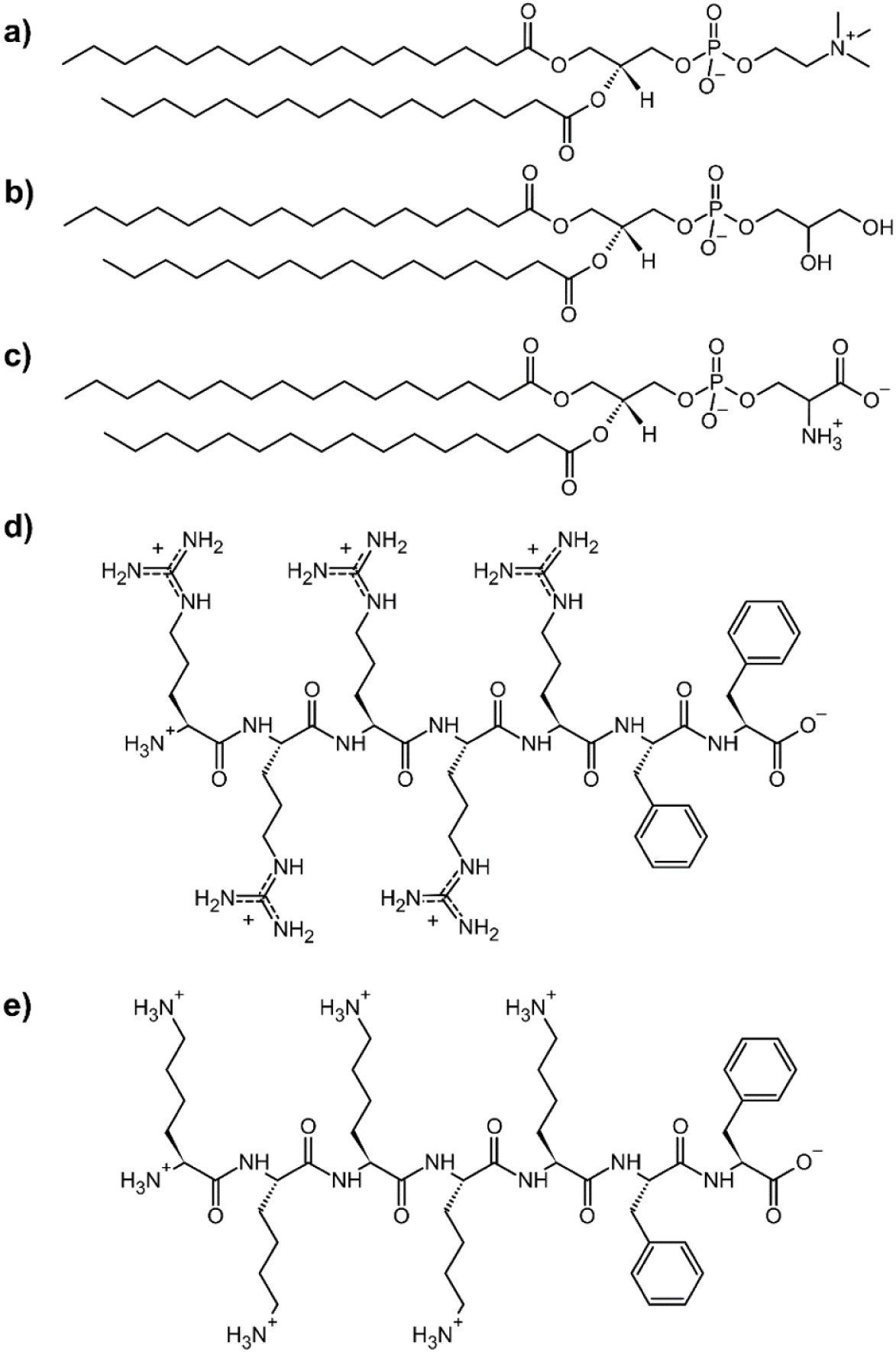
Structural formulas of: a) 1,2-dipalmitoyl-*sn*-glycero-3-phosphocholine (DPPC); b) 1,2-dipalmitoyl-*sn*-glycero-3-phospho-(1′-rac-glycerol) sodium salt (DPPG); c) 1,2-dipalmitoyl-*sn*- glycero-3-phosphoserine (DPPS); d) R5F2; e) K5F2.

**Fig. 2.**
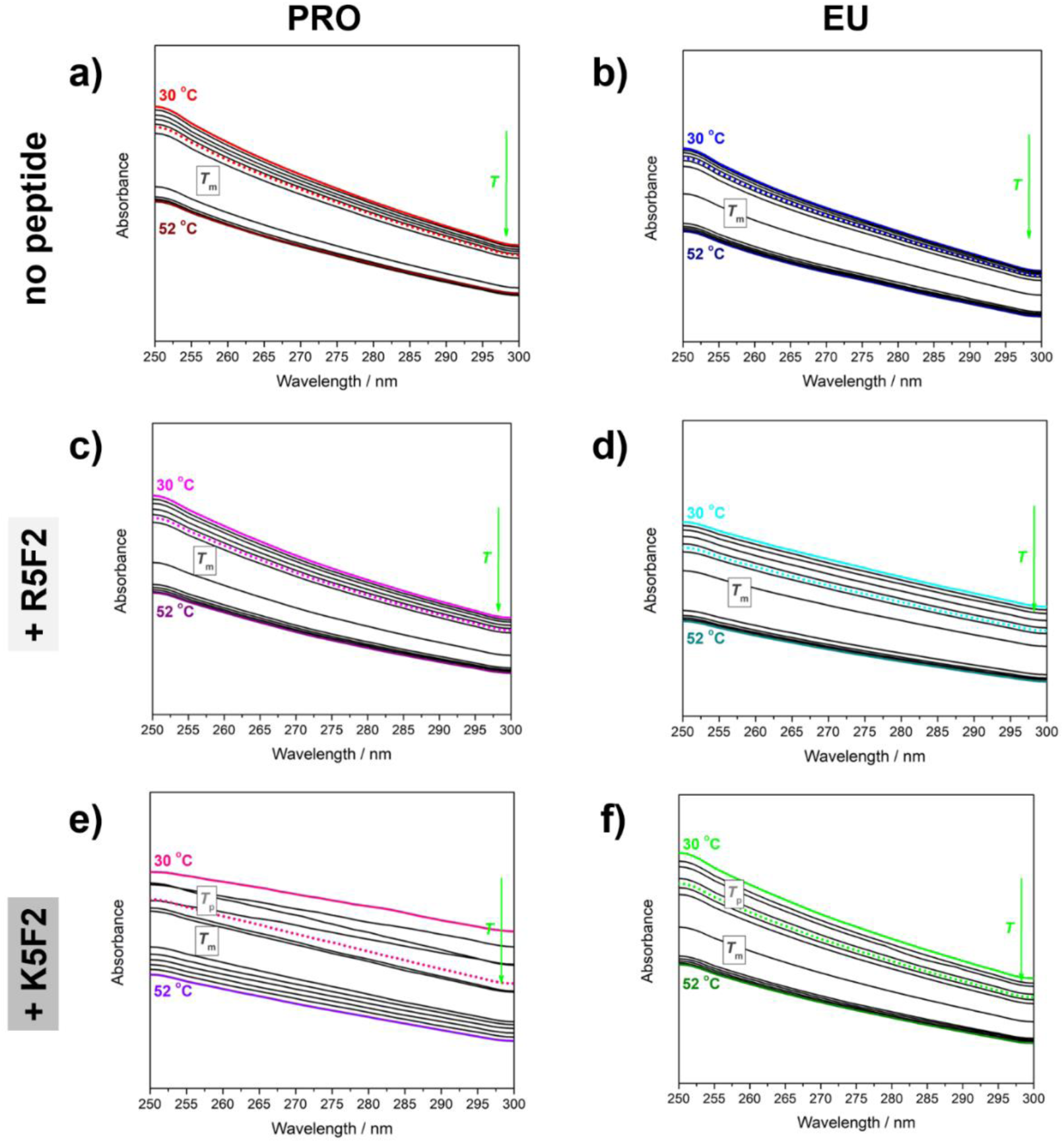
Temperature-dependent UV-Vis spectra (solid curves) and spectral profiles (dotted curves) of: a) PRO; b) EU; c) PRO+R5F2; d) EU+R5F2; e) PRO+K5F2; f) EU+K5F2. The spectra acquired at 30 °C/52 °C are highlighted (red/wine for PRO, blue/navy for EU, magenta/purple for PRO+R5F2, cyan/dark cyan for EU+R5F2, pink/violet for PRO+K5F2, green/olive for EU+K5F2), as well as spectral profile (red for PRO, blue for EU, magenta for PRO+R5F2, cyan for EU+R5F2, pink for PRO+K5F2, green for EU+K5F2).

As the details on MCA are thoroughly explained in other papers [33,45–47], only the fundamentals will be outlined in the continuation of the text. The collected spectra can be presented in a matrix form (**D**) as the product of two matrices: the one that presents the concentrational profile (**C**) of one component that exhausts all temperature-dependent variability in the spectra, and the other that represents its spectral profile (**S**):

**D**=**CS^T^**+**E**,

where **E** presents matrix of residuals unexplained by the **CS^T^** product.

The obtained concentrational profile of this one component displays (a single or a double) sigmoidal character in all six examined systems (PRO ± R5F2/K5F2 and EU ± R5F2/K5F2) and the obtained inflection points perfectly coincide with the phase transition temperatures (*T*_pt_), both *T*_p_ and *T*_m_ [41,43]. By fitting the concentration profile on a single/double Boltzmann fit the following *T*_pt_ values were obtained: PRO: *T*_m_ = 41.5 ± 0.3 °C (*R*^2^=0.999); PRO+R5F2: *T*_m_ = 39.5 ± 0.4 °C (*R*^2^=0.999); PRO+K5F2: *T*_p_ = 35 ± 1 °C (*R*^2^=0.999); EU: *T*_m_ = 41.7 ± 0.4 °C (*R*^2^=0.998); EU+R5F2: *T*_m_ = 41.0 ± 0.3 °C (*R*^2^=0.971); EU+K5F2: *T*_p_ = 34 ± 2 °C, *T*_m_ = 41.2 ± 0.4 °C (*R*^2^=0.998).

### 2.5 FTIR-ATR Spectroscopy of LUVs: Data Acquisition and Spectral Analysis

Invenio-S Bruker spectrometer equipped with the photovoltaic LN-MCT detector and BioATR unit was used for measurements of FTIR spectra of PRO ± R5F2/K5F2 and EU ± R5F2/K5F2 suspensions (*γ*(lipids)= 5 mg mL^−1^), as well as of and R5F2/K5F2 (*γ*(R5F2/K5F2) = 1 mg mL^-1^) and PB solutions (peptide spectra are presented in Supporting Information, Fig. S3). The circular BIOATR II unit (radius of 2 mm) is based on dual crystal technology, where the upper ATR crystal is made of silicon and the lower ATR crystal is made of ZnSe. The inside of the ATR unit was continuously purged with N_2_ gas connected with an external supply and temperature-controlled using a circulating water bath of Huber Ministat 125. The solutions of R5F2/K5F2 and suspensions PRO ± R5F2/K5F2 and EU ± R5F2 were pipetted directly on the ATR crystal unit (30 μL of solution/suspension) and their spectra were acquired against air as a background. For suspensions, at least three independent unit fillings were taken and one for reference solution. In all measurements, the air was used as background. All spectra were collected with a nominal resolution of 2 cm^-1^ and 256 scans using OPUS 8.5 SPI (20200710) software.

Following the subtraction of PB spectrum from PRO ± R5F2/K5F2 and EU ± R5F2/K5F2 spectra acquired at the same temperatures, the obtained difference FTIR spectra were examined in the following spectral regions [42]: i) 2980-2820 cm^−1^, ii) 1700-1690 cm^−1^, iii) 1515-1395 cm^−1^, iv) 1395-1325 cm^−1^; v) 1275-1130 cm^−1^ and vi) 1130-995 cm^−1^. In the listed spectral regions one can observe the bands originated from: i) (anti)symmetric stretching of methylene moieties (ν_(a)s_CH_2_) of hydrocarbon chains; ii) carbonyl stretching of glycerol backbone that can be either non-hydrogen-bonded (non-HB) or hydrogen-bonded (HB) (νC=O_(non-)HB_), iii) scissoring of methylene groups (γCH_2_), iv) scissoring of methyl groups of hydrocarbon chains and wagging of methylene groups (γCH_3_, ωCH_2_), v) antisymmetric stretching of phosphate groups and glycerol moieties (ν_as_PO_2_^−^ and ν_(a)s_C−O) and vi) symmetric stretching of phosphate and their neighboring C−O groups (ν_s_PO_2_^−^ and νCOP) along with antisymmetric stretching of choline moiety (ν_as_C−N) [48–50]. In the selected regions the spectra were smoothed (Savitzky-Golay; polynomial of a 3^rd^ degree through 30 points), baseline corrected (two points) and normalized [42].

### 2.6 Molecular dynamics simulations

Classical molecular dynamics (MD) was utilized for modeling PRO and EU bilayers and their interactions with R5F2 or K5F2. The bilayers were prepared by CHARMM-GUI [51]. PRO bilayer consisted of 230 DPPC lipids and 26 DPPG lipids, while EU bilayer consisted of 230 DPPC lipids and 26 DPPS lipids. The bilayers were solvated with 75 waters per lipid molecule, and NaCl was added to neutralize and achieve the concentration of 100 mmol dm^−3^. 8 peptides (either R5F2 or K5F2) were then randomly inserted into the water phase. The systems were minimized and heated for 200 ps in the NVT ensemble using the V-rescale algorithm. The production was run in the NpT ensemble. The Nosé-Hoover thermostat maintained the temperature at either 30 or 50 °C, with a time constant of 1 ps. The Parrinello-Rahman barostat maintained the pressure of 1 bar with the semi-isotropic scaling and time constant of 5 ps. The total production time was 500 ns. Analyses such as structural parameters or density profiles were conducted only on the last 300 ns of production, with the initial segment dismissed as equilibration time.

The simulations were run in GROMACS 2020.0 software [52]. Lipids and peptides were described by the CHARMM36m force field [53], and the TIP3 model was used for water [54]. The cutoff for short-range Coulomb interactions and van der Waals interactions was 1.2 nm with a switching function turned on after 1.0 nm, and long-range Coulomb interactions were handled with Particle mesh Ewald (PME). The bonds involving hydrogen were constrained using LINCS, and the time step was 2 fs. Three-dimensional periodic boundary conditions were applied throughout.

The simulations were conducted on the Supek supercomputer at the University Computing Centre (SRCE) in Zagreb, Croatia [55].

### 2.7 Confocal microscopy of GUVs

Imaging was performed on a Leica SP8 confocal setup (Mannheim, Germany). The fluorescent dye, DilC_18_, was excited with a 552 nm laser. Before imaging samples, cover glasses used to prepare the observation chambers were coated via incubation with 10 mg/ml bovine serum albumin (BSA, Sigma) in pure water solution to prevent vesicle adhesion. Before adding the vesicle suspension, the glasses were rinsed with pure water to remove any unbound BSA. GUVs of two different compositions were measured, POPC+10% POPG (PRO’) and POPC+10% POPS (EU’) in the absence/presence of R5F2 and K5F2. For imaging, 15 µL of the desired GUVs suspension was placed on BSA-coated glass and diluted with 45 µL of glucose solution (same osmolarity as that of the sucrose solution), or with a 1:1 ratio solution using 10 µL of the GUV suspension and 10 µL of glucose solution. All GUV samples were first observed without peptides and then in addition of 1 µM, 5 µM, 10 µM or 20 µM peptides. All images were analyzed using ImageJ software.

## 3. Results

### 3.1. Thermotropic properties of PRO/EU systems ± R5F2/K5F2: DSC and UV-Vis data

In general, in the absence of peptides PRO and EU systems display pretransition (*T*_p_) only as a barely seen deviation from the baseline, while the main phase transition (*T*_m_) appears as a broad envelope with one (EU) or two unresolved (PRO) maxima (Fig. 3a and b). On the other hand, the concentrational profile of projected temperature-dependent UV-Vis spectra appears as a sigmoidal curve having only one inflection point. The obtained values for PRO system obtained from DSC/UV-Vis are *T_p_ ∼* 33. 2 °C, *T_m,1_* = 40.0 ± 0.1 °C and *T_m,2_* = 40.6 ± 0.1 °C (DSC)/*T_m_* = 41.5 ± 0.3 °C (UV-Vis), whereas for the analogous EU system the corresponding values are *T_p_ ∼* 37 °C and *T_m,1_* = 41.6 ± 0.1 °C (DSC)/*T_m_* = 41.7 ± 0.4 °C (UV-Vis) (Table 1). Since the resulting *T*_p_ in both PRO and EU systems in the absence of peptides is only an estimate, the study of eventual variations is trivial in the scope of this paper. As far as the main phase transition of PRO system in the DSC experiment is concerned, it is composed of two processes separated by about 0.6 °C, presumably due to the change in the curvature and bilayer thinning [41]. Moreover, *T*_m,1/2_ values of PRO differ for about 1 °C in comparison with *T*_m,1_ measured in EU system (41.6 ± 0.1 °C), which suggests that in EU system the gel phase is more stabilized than in PRO systems [36,56,57]. The values obtained from UV-Vis measurements of both PRO and EU systems are practically the same, implying that from the turbidity aspect, there are no differences between the two of them.

**Fig. 3.**
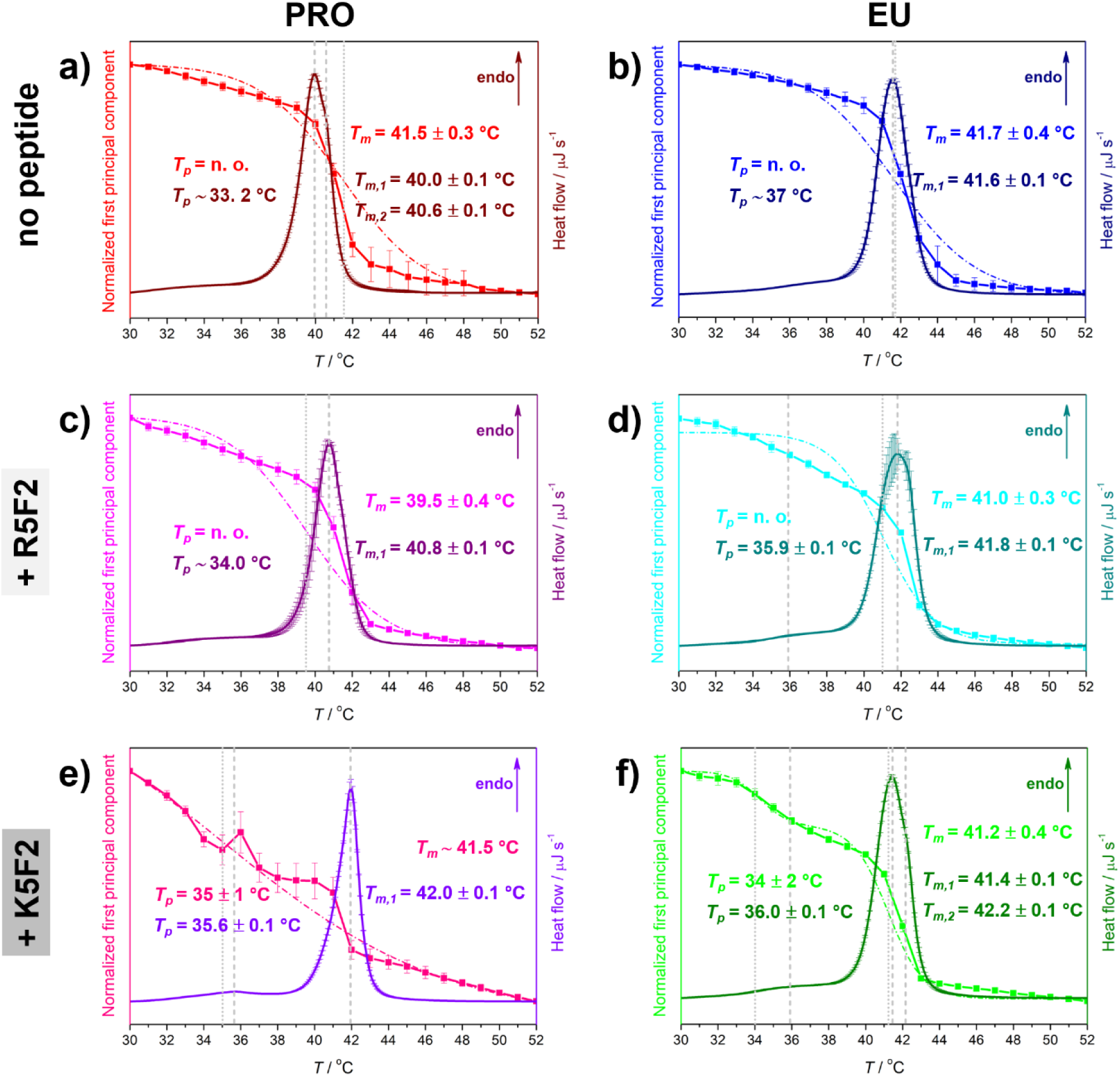
DSC curves and concentrational profiles of the (first) principal component accompanied with a single or a double Boltzmann sigmoidal transition of: a) PRO system in the absence of peptides (wine curve for DSC and red curve for spectral projection of UV-Vis data (solid)/double Boltzmann fit (dash-dot curve)); b) EU system in the absence of peptides (navy curve for DSC and blue curve for spectral projection of UV-Vis data (solid)/double Boltzmann fit (dash-dot curve)); c) PRO+R5F2 system (purple curve for DSC and magenta curve for spectral projection of UV-Vis data (solid)/double Boltzmann fit (dash-dot curve)); d) EU+R5F2 system (dark cyan curve for DSC and cyan curve for spectral projection of UV-Vis data (solid)/double Boltzmann fit (dash-dot curve)); e) PRO+K5F2 system (violet curve for DSC and pink curve for spectral projection of UV-Vis data (solid)/double Boltzmann fit (dash-dot curve)); f) EU+R5F2 system (olive curve for DSC and green curve for spectral projection of UV-Vis data (solid)/double Boltzmann fit (dash-dot curve)). Phase transition temperatures are highlighted with dashed (DSC) and dotted (UV-Vis) lines and are additionally written on graphs and designated with a corresponding color.

**Table 1.** Temperatures of the phase transitions (*T*_pt_) of PRO/EU systems in the absence/presence of R5F2/K5F2 peptides determined from the maxima of DSC curves (*T*_p_ and *T*_m, 1/2_ for pre-and the main phase transition, respectively) and from inflection points of concentration profiles obtained from multivariate curve analysis (MCA; *T*_p_ and *T*_m_ for pre-and the main phase transition, respectively) by estimation/fitting on a single/double Boltzmann profile.

| system | $T_{pt}^a$ | | | |
| --- | --- | --- | --- | --- |
|  | DSC |  | UV-Vis |  |
| | $T_p$ | $T_{m, 1/2}$ | $T_p$ | $T_m$ |
| PRO | $\sim 33.2$ | $40.0 \pm 0.1$ | n. o. | $41.5 \pm 0.3$ |
| | | $40.6 \pm 0.1$ | | |
| PRO+R5F2 | $\sim 34$ | $40.8 \pm 0.1$ | n. o. | $39.5 \pm 0.4$ |
| PRO+K5F2 | $35.6 \pm 0.1$ | $42.0 \pm 0.1$ | $35 \pm 1$ | $\sim 41.5$ |
| EU | $\sim 37$ | $41.6 \pm 0.1$ | n. o. | $41.7 \pm 0.4$ |
| EU+R5F2 | $35.9 \pm 0.1$ | $41.8 \pm 0.1$ | n. o. | $41.0 \pm 0.3$ |
| EU+K5F2 | $36.0 \pm 0.1$ | $41.4 \pm 0.1$ | $34 \pm 2$ | $41.2 \pm 0.4$ |
| | | $42.2 \pm 0.1$ | | |

<sup>a</sup> In °C; n. o. stands for not observed; ∼ refers to the estimation of the baseline deviation

In the presence of R5F2, the DSC curve of the PRO system provides only a rough estimation of the pretransition and the main phase transition characterized by one maximum only, whereas the corresponding UV-Vis data displays a concentration profile of spectral projection having one inflection point;i.e. the measured (DSC)/obtained (UV-Vis) values are: *T_p_ ∼* 34.0 °C and *T_m,1_* = 40.8 ± 0.1 °C (DSC)/*T_m_* = 39.5 ± 0.4 °C (UV-Vis) (Fig. 3c and d). Interestingly, from the DSC curve of EU system in the presence of R5F2 it became possible to estimate a minor peak at the expected pretransition temperature and a broad signal associated with the main phase transition of EU-composed LUVs (*T_p_ =* 35.9 ± 0.1 °C and *T_m,1_* = 41.8 ± 0.1 °C), while UV-Vis measurements reported the discontinuous turbidity change at the value that approximately coincides with the main phase transition temperature determined from DSC (*T_m_* = 41.0 ± 0.3 °C). As in the peptide-free systems, the *T*_m_ values of the EU system are higher than the corresponding values in the PRO system. However, in the presence of R5F2, the difference is reduced at the expense of an increase in *T*_m_ of the PRO systems (Table 1).

In contrast to the discussed four systems (PRO/EU ± R5F2), the presence of K5F2 in PRO and EU systems induces stronger pretransition as observed in both DSC and UV-Vis data (Fig. 3e and f). Moreover, the main phase transition itself exhibits clear differences: in the PRO system, the *T*_m_ determination from UV-Vis data is obtained only by visual estimation of the inflection point, whereas in the EU system the main phase transition is displayed as two unresolved maxima in DSC curve. Accordingly, the DSC/UV-Vis values of/in the PRO+K5F2 system are as follows: *T_p_ =* 35.6 ± 0.1 °C and *T_m,1_* = 42.0 ± 0.1 °C (DSC)/*T_p_* = 35 ± 1 °C and *T_m_* ∼ 41.5 °C (UV-Vis), while the analogous measured (DSC)/obtained (UV-Vis) values for EU+K5F2 system are *T_p_ =* 35.6 ± 0.1 °C and *T_m,1_* = 42.0 ± 0.1 °C (DSC)/*T_p_* = 34 ± 2 °C and *T_m_* = 41.2 ± 0.4 °C (UV-Vis). The similarity between the values measured from DSC experiments, as well as those determined (and estimated) from UV-Vis measurements (Table 1), suggest that the presence of K5F2 peptide impacts PRO and EU lipid bilayers twofold: i) it amplifies the ripples at the surfaces of both PRO and EU lipid bilayers at temperatures below *T*_m_ and ii) makes both systems more rigid (larger *T*_m_) in comparison with pure PRO and EU lipid bilayers. Notably, the turbidity-based response (UV-Vis) suggests that the pre-and the main phase transition in PRO+K5F2 are coupled, which contrasts the response of EU+K5F2 system in which these two thermotropic events are well separated. The observed phenomenon may be associated with both the affinity and the dynamics of K5F2 adsorption/desorption on the surface of lipid bilayers [23].

### 3.2. Molecular properties of PRO/EU systems ± R5F2/K5F2 as revealed from FTIR data

The analysis of the first spectral region of FTIR spectra (2980-2820 cm^−1^) in all six systems (PRO/EU ± R5F2/K5F2) does not reveal any particular surprises; owing to the weakening of van der Waals interactions between the hydrocarbon chains of lipid molecules upon heating, the band originated from ν_as_CH_2_ displaces from 2919 cm^−1^ (30 °C) to 2924 cm^−1^ (50 °C), and the one from ν_s_CH_2_ displaces from 2851 cm^−1^ (30 °C) to 2854 cm^−1^ (50 °C) (Fig. 4a and d) [23,48].

**Fig. 4.**
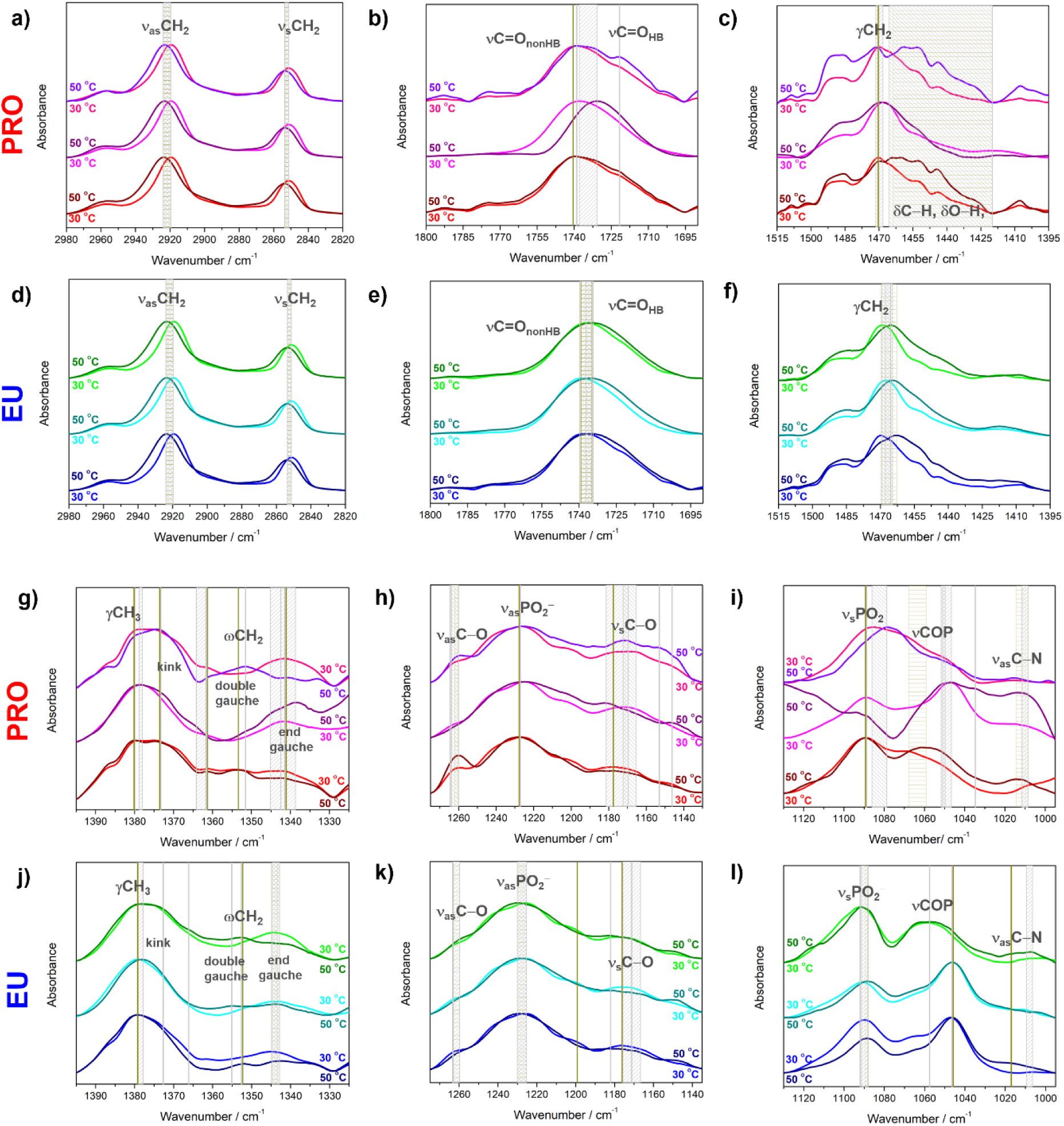
Normalized, smoothed and baseline-corrected FTIR spectra of PRO ± R5F2/K5F2 and EU ± R5F2/K5F2 in the following spectral ranges: a, d) 2980-2820 cm^−1^ (ν_(a)s_CH_2_); b, e) 1700-1690 cm^−1^ (νC=O_non-)HB_); c, f) 1515-1395 cm^−1^(γCH_2_, δC−H, δO−H); g, j) 1395-1325 cm^−1^ (γCH_3_, ωCH_2_); h, k) 1275-1130 cm^−1^ (ν_(a)s_C−O, ν_(a)s_PO ^−^); i, l) 1130-995 cm^−1^ (ν_s_PO ^−^, νCOP, ν_as_C−N). PRO, PRO+R5F2 and PRO+K5F2 spectra at 30 °C/50 °C are presented with solid red/wine, magenta/purple, and pink/violet curves, respectively; EU, EU+R5F2 and EU+K5F2 spectra at 30 °C/50 °C are presented with solid blue/navy, cyan/dark cyan, and green/olive curves, respectively. Along with the band assignment, their displacements in the presence of peptides are labeled with light gray-shaded rectangles, while in their absence with dark yellow, respectively. The bands that maintain the position upon the gel-to-fluid phase transition are designated with solid light gray (in the presence of peptides) or dark yellow (in the absence of peptides) lines.

The most important band that appears in the second spectral region (1800-1690 cm^−1^) emerges from the stretching of carbonyl group (νC=O) of glycerol backbone that is (not) engaged in a hydrogen bond formation with surrounding water molecules and other hydrogen bond donating and accepting centers (C=O_non-HB_ and C=O_HB_) [58–60]. In PRO system, in the absence of peptides, the maximum of the band does not change upon the phase transition, i.e. the band maximum remains at 1741 cm^−1^ (at 30 °C and 50 °C), but the change in the shape, emphasizing the asymmetry on the low-frequency side, is linked with an increase of C=O_HB_ population. In PRO+R5F2 the heating induces low-frequency shift of this broad band from 1738 cm^−1^ (30 °C) to 1731 cm^−1^ (50 °C), suggesting the increase of the C=O_HB_ population the expense of C=O_non-HB_ [59,61,62]. Upon the gel-to-fluid phase transition of PRO+K5F2 system, this increase in the population of C=O_HB_ is even more pronounced since the broad band having a maximum at 1739 cm^−1^ (30 °C) is maintained and accompanied by the additional νC=O_HB_ band with distinguished maximum at 1722 cm^−1^ (50 °C). This peptide-dependent response of PRO system suggests the diversity in their interaction with glycerol moiety, highlighting the substantial rise of C=O_HB_ population in the presence of K5F2, that ultimately reflects both variations in the bilayer penetration depth and the strength of the adsorption (Fig. 4b). In EU systems, the overall band shape generated by C=O groups, as well as the band displacement upon gel-to-fluid transition, is remarkably similar regardless of the absence or presence of peptides (Fig. 4e). In particular, in the absence of peptides the maximum of a broad envelope displaces from 1739 cm^−1^ (30 °C) to 1735 cm^−1^ (50 °C), whereas in the presence of R5F2 and K5F2 the corresponding shift is observed from 1740 cm^−1^ (30 °C) to 1735 cm^−1^ (50 °C) and from 1737 cm^−1^ (30 °C) to 1735 cm^−1^ (50 °C), respectively. The only truly measured difference between peptide-dependent systems is the position of the band maximum in the gel phase (at 30 °C; R5F2/K5F2 produces the maximum at 1740 cm^−1^/1737 cm^−1^), suggesting that the presence of R5F2 may reduce the hydration of glycerol moiety and/or simply prevents its involvement in a hydrogen bond network as in the case of K5F2 (Fig. 4e).

A third examined spectral region (1515-1395 cm^−1^) is distinguished by the signals originating from the scissoring of methylene groups of hydrocarbon chains that are exceptionally sensitive to lateral interactions between lipid molecules [48,63,64]. In PRO system in the absence of peptides in the gel phase the band maximum appears at 1470 cm^−1^ (30 °C) and remains at the same position; the same behavior is observed for the fluid phase (at 1470 cm^−1^ at 50 °C) (Fig. 4c), indicating the packing of PRO system lipids in presumably orthorhombic arrangement [48,65]. The appearance of several maxima in a gray-shaded region immediately below 1470 cm^−1^, likely assigned as deformations of −CH and −OH moieties of DPPG [66], may camouflage the existence of an additional maximum at 1465 cm^−1^, which is anticipated for the lipid ordering in an orthorhombic arrangement. In the presence of R5F2, the displacement of the band maximum from 1468 cm^−1^ (30 °C) to 1470 cm^−1^ (50 °C) suggest a change in the packing pattern from hexagonal to a disordered fluid [65]. On the contrary, K5F2 induces a very small displacement of the corresponding band maximum to the opposite direction upon heating, i.e. from 1470 cm^−1^ (30 °C) to 1471 cm^−1^ (50 °C), suggesting the possible maintenance of the preferred orthorhombic packing pattern of lipid molecules [67]. Additionally, unlike R5F2, the presence of K5F2 preserves the structure of a broad envelope that encompasses the region ∼ 1470-1420 cm^−1^ (a gray-shaded region in Fig. 4c), very likely resulting from a different interaction pattern of these peptides with PRO system. In EU-related systems the spectral assignment seems to be a bit simpler than in analogous PRO systems as in both absence and the presence of peptides there is a low-frequency shift of the band maximum upon gel-to-fluid phase transition of lipid bilayers (Fig. 4f). In particular, in the peptide-free EU system, the band maximum displaces from 1470 cm^−1^ (30 °C) to 1463 cm^−1^ (50 °C), in EU+R5F2 from 1468 cm^−1^ (30 °C) to 1465 cm^−1^ (50 °C), and in EU+K5F2 from 1469 cm^−1^ (30 °C) to 1465 cm^−1^ (50 °C). The magnitude of the displacement suggests that the hexagonal arrangement in the EUs system in the gel [48] transforms to disordered arrangement in fluid phase.

The most distinguished signals in the fourth spectral region (1395-1325 cm^−1^) are those originated from the umbrella of methyl groups of hydrocarbon chains (γCH_3_) and a series of bands attributed to the wagging of hydrocarbon chains methylene groups (ωCH_2_), whose intensity and position can be associated with the *gauche* conformers of certain type, as well as with the existence of conformers that exhibit a kink [63,64,68]. Despite the complexity of the observed envelope, it is clear that the γCH_3_ band in PRO and EU systems in the absence of peptides does not displace upon the gel-to-fluid phase transition: in the former it appears at 1380 cm^−1^ (at 30 °C and 50 °C) and in the latter at 1379 cm^−1^ (Fig. 4g and j). In the presence of R5F2, both PRO and EU lipid bilayers exhibit a very small shift of the γCH_3_ band; in PRO+R5F2 it displaces from 1378 cm^−1^ (30 °C) to 1379 cm^−1^ (50 °C), while the opposite direction is observed for EU+R5F2, i.e. from 1379 cm^−1^ (30 °C) to 1378 cm^−1^ (50 °C). Upon the adsorption of K5F2, an analogous phenomenon as for peptide-free PRO and EU systems is observed, except for a wavenumber maximum position: in PRO+K5F2 the band remains at 1380 cm^−1^ (at 30 °C and 50 °C), while in EU+K5F2 at 1379 cm^−1^ (at 30 °C and 50 °C). A more complicated phase-induced response is registered regarding the signals assigned to the ωCH_2_ bands, especially in PRO system (Fig. 4g). First, in the absence of peptides the band maxima attributed to the wagging of kink conformers are detected at 1373 cm^−1^ (at 30 °C and 50 °C) and at 1361 cm^−1^ (at 30 °C and 50 °C), those assigned as the wagging of double *gauche* conformers at 1354 cm^−1^ (at 30 °C and 50 °C) and the band maximum attributed to end *gauche* conformers at 1342 cm^−1^ (at 30 °C and 50 °C). In PRO+R5F2 the bands due to the wagging of kink conformers appear at 1373 cm^−1^ (at both 30 °C and 50 °C) and at 1364 cm^−1^ (at 30 °C) that displaces to 1361 cm^−1^ (50 °C), the double *gauche* conformers generate the band with maximum at 1352 cm^−1^ (at 30 °C and 50 °C), whereas the signal originated from end *gauche* conformers at 1342 cm^−1^ (30 °C) upon gel-to-fluid phase transition extends from ∼ 1345-1338 cm^−1^ (50 °C). The analogous signals in PRO+K5F2 display maxima at 1374 cm^−1^ (at both 30 °C and 50 °C) and 1361 cm^−1^ (30 °C and 50 °C) (kink conformers), at 1352 cm^−1^ (only at 50 °C) (double *gauche*) and at 1341 cm^−1^ (at 30 °C and 50 °C) (end *gauche*). A major difference across PRO systems is that the presence of K5F2 eliminates (or, at best, suppresses) the signal of double *gauche* conformers at 30 °C, but then its signal rises at the expense of the reduction of the intensity of the band assigned as the wagging of end *gauche* groups. The inversion of end *gauche* and double *gauche* band intensities is not unusual during melting [64], but it is rather interesting that it is more pronounced for PRO+K5F2 than in any other system. In EU systems in the absence of peptides the band maxima of double *gauche* conformers are found at 1352 cm^−1^ (at 30 °C and 50 °C) and those of end *gauche* conformers are displaced from 1345 cm^−1^ (30 °C) to 1343 cm^−1^ (50 °C). In EU+R5F2 the band maxima associated with the wagging of double *gauche* conformers are found at 1355 cm^−1^ (30 °C and 50 °C), while the maximum of the end *gauche* conformers is displaced from 1345 cm^−1^ (30 °C) to 1342 cm^−1^ (50 °C). The gel-to-fluid phase transition is accompanied by the redistribution of the intensities of these two bands, i.e. the latter expectedly increases at the expense of the former upon melting [48] (Fig. 4j). Aside from the detectable maxima at 1373 cm^−1^ (at 50 °C) and 1366 cm^−1^ (at 50 °C), which originate from the resolved kink conformer, the signals of double *gauche* and end *gauche* conformers in EU+K5F2 resemble those of EU+R5F; double *gauche* conformers produce the signal at 1352 cm^−1^ (at 30 °C and 50 °C), whereas end *gauche* ones, with a maximum at 1345 cm^−1^ (30 °C) displaced to 1342 cm^−1^ (50 °C), are accompanied with the inversion of the band intensity owing to an increase of a double *gauche* conformers upon melting [64].

The fifth spectral region (1275-1130 cm^−1^) is the most distinguished by the signals originating from antisymmetric stretching of phosphate groups (ν_as_PO_2_^−^) of lipid molecules that, as well as C=O moieties, can be non-HB (ν_as_PO_2_^−^_nonHB_) and HB (ν_as_PO_2_^−^_HB_) [50]. In PRO systems, irrespective to the absence/presence of peptides, the broad envelope generated by both ν_as_PO_2_^−^_nonHB/HB_ displays a maximum at the same wavenumber that does not change upon the gel-to-fluid phase transition, i.e. in PRO ± R5F2/K5F2 it appears at 1227 cm^−1^ (at 30 °C and at 50 °C) (Fig. 4h). The corresponding band in all EU systems exerts the same response, and, unlike in PRO systems, there is an obvious shift of the band maximum, i.e. there is a high-frequency displacement upon gel-to fluid phase transition from 1225 cm^−1^ (30 °C) to 1231 cm^−1^ (50 °C) (Fig. 4k). As the observed band is a superposition of numerous bands generated by phosphate groups in different surroundings roughly divided into two subpopulations (non-HB and HB) [69], from the observed phenomena one can conclude that the phase transition causes the increase in the number of non-HB phosphate groups in EU systems, i.e. very likely their decrease hydration. On the other hand, besides small qualitative changes in the envelope shape, in PRO systems nothing drastically changes upon the phase transition, regardless of the peptides’ presence, so the peptides do not appear to severely affect the hydrogen bond network meshed by phosphate groups. Along with the signals of phosphate groups, in the examined spectral region the signals originated from the (anti)symmetric stretching of C-O moieties (ν_(a)s_C−O) appear as well [50]; in PRO ν_as_C−O and ν_s_C−O appear at 1261 cm^−1^/1260 cm^−1^ (30 °C/50 °C) and at 1178 cm^−1^/1178 cm^−1^ (30 °C/50 °C), in PRO+R5F2 at 1264 cm^−1^/− (30 °C/50 °C) and at 1170 cm^−1^/1182 cm^−1^ (30 °C/50 °C), and in PRO+K5F2 at 1264 cm^−1^/1260 cm^−1^ (30 °C/50 °C) and at 1166 cm^−1^/1172 cm^−1^ (30 °C/50 °C), respectively (Fig. 4h). The corresponding signals in the EU system display maxima at 1263 cm^−1^/1260 cm^−1^ (30 °C/50 °C) and at 1176 cm^−1^/1176 cm^−1^ (30 °C/50 °C); in EU+R5F2 at 1263 cm^−1^/− (30 °C/50 °C) and at 1176 cm^−1^/1166 cm^−1^ (30 °C/50 °C), and in EU+K5F2 at 1263 cm^−1^/1260 cm^−1^ (30 °C/50 °C) and at 1173 cm^−1^/1182 cm^−1^ (30 °C/50 °C), respectively (Fig. 4k). Additionally, in all systems weak bands appear at between 1160-1140 cm^−1^, that most likely originate from various deformation modes of C−H groups [64].

In the sixth spectral region examined (1130-975 cm^−1^), the most important lipid-originated bands are those that arise due to the symmetric stretching of phosphate groups (ν_s_PO_2_^−^) and due to the stretching of P−O−C− lipid parts (νPOC); the former usually appears at about 1080-1090 cm^−1^, the latter between about 1050-1070 cm^−1^, and, even though both envelopes are rather strong, the former usually exceeds the latter in the intensity (Fig. 4i and l) [15,49]. Upon gel-to-fluid phase transition in both systems in the absence of peptides ν_s_PO_2_^−^ band remains at the same position. In PRO it appears at 1090 cm^−1^/1090 cm^−1^ (30 °C/50 °C), and in EU at 1092 cm^−1^/1092 cm^−1^ (30 °C/50 °C). The νPOC band either displaces to the lower frequencies (PRO) from 1068 cm^−1^ (30 °C) to 1059 cm^−1^ (50 °C), or remains at the same position (EU), i.e. it appears at 1057 cm^−1^ /1057 cm^−1^ (30 °C/50 °C). Along with these bands there are choline group-associated bands (ν_as_CN) [50] displaced from 1011 cm^−1^ (30 °C) to 1013 cm^−1^ (50 °C) in PRO and at 1017 cm^−1^ (only at 50 °C) in EU systems in peptide-free conditions. In the presence of R5F2 in PRO system ν_s_PO_2_^−^ band maintains its position maximum at 1089 cm^−1^ (at 30 °C and 50 °C), whereas in EU it slightly displaces to lower frequencies, i.e. from 1091 cm^−1^ (30 °C) to 1088 cm^−1^ (50 °C). The inversion of band maximum displacement is seen for the νPOC band: in PRO it displaces from 1047 cm^−1^ (30 °C) to 1050 cm^−1^ (50 °C), while in EU the position of the corresponding band remains at 1046 cm^−1^ (at 30 °C and 50 °C). The features of ν_as_CN are significantly enhanced in PRO system in the presence of R5F2; upon gel-to-fluid phase transition one additional maxima appears at 1035 cm^−1^ (50 °C) and a remarkable intensity gain of the band displaced from 1010 cm^−1^ (30 °C) to 1012 cm^−1^ (50 °C) band is detected. In EU+R5F2 system there is no analogous or comparable effect. Ultimately, in PRO+K5F2 system ν_s_PO_2_^−^ and νPOC band maxima displace from 1086 cm^−1^ (30 °C) to 1079 cm^−1^ (50 °C) and from 1050 cm^−1^ (30 °C) to 1052 cm^−1^ (50 °C), respectively, whereas in EU+K5F2 they position maxima remain at the same wavenumber; at 1092 cm^−1^ (at 30 °C and 50 °C) and at 1057 cm^−1^ (at 30 °C and 50 °C), respectively. Additionally, ν_as_CN signals are detectable only in EU+K5F2 as very weak bands at 1010 cm^−1^ (30 °C) and 1007 cm^−1^ (50 °C). As one of the most significant features associated with two of the most distinguished bands is addressed to their intensities ratios; usually, the bands are either of comparable intensities or the ν_s_PO_2_^−^ band is slightly stronger than νPOC. In PRO systems in the presence of R5F2 the expected intensity ratio becomes significantly inverted simultaneously with the appearance of, very likely, another band at about 1035 cm^−1^ that gain intensity upon gel-to-fluid phase transition (Fig. 4i.) Interestingly, in PRO+K5F2 these bands are considerably superimposed and the former is by far stronger than the latter, regardless of the phase of the lipid bilayers. In EU and EU+R5F2 systems the latter band is slightly stronger than the former, but in EU+K5F2 the former is enhanced and the latter is broader, irrespective to the gel/fluid phase (Fig. 4l). Although these bands are usually addressed as not as useful in diagnostic purposes [48], their intensity ratios imply that transitional dipole moments associated with normal modes are changed in the presence of R5F2/K5F2, as well as the fact that 10 % of DPPG/DPPS (mole fraction) in dominantly DPPC bilayer also affects the response.

The differences in the responses of PRO and EU systems, irrespective of the presence of peptides, may occur due to the hydration pattern of lipid bilayers that differ in the type of anionic lipids at a mole fraction of 10 %. Additionally, these differences may arise from the peptides that inherently form different kinds of intermolecular interactions under the givenposed conditions. The contribution of these factors was evaluated with the aid of MD simulations.

### 3.3 Molecular dynamics simulations results

MD simulations were employed to investigate the interactions of peptides from the aqueous medium with the bilayers. Basic structural parameters used to confirm the correct phase of a bilayer are reported in Supporting Information, Section S5 (Figs. S4 and S5). The adsorption was tracked by calculating the minimum distance of any peptide atom to the P-atom, while the preferred location and orientation are visible from number density profiles. Peptides were shown to primarily interact with the lipid interface through the charged amino acids, but the propensity for binding varied between systems (Figs. 5 and S5, 30 °C data separated to Supporting Information to improve legibility). PRO bilayers adsorbed R5F2 quickly and strongly, particularly in the fluid phase, and after 300 ns all peptides were bound and remained on the surface for the duration of the simulation. In contrast, K5F2 was less inclined to bind the bilayer and more likely to dissociate during the simulation. R/K residues were retained among the headgroups, with F mainly facing the solvent, but in the 50 °C simulation, there were also instances of peptide penetration into the hydrophobic bilayer environment. EU bilayers showed similar behavior, with R5F2 binding within the first 100 ns, though at 30 °C there was an instance of dissociation. K5F2 peptides were weakly interacting with the surface with a lot of dissociation events. Insertion still occurred at 50 °C for both peptides, but also at 30 °C for R5F2.

**Fig. 5.**
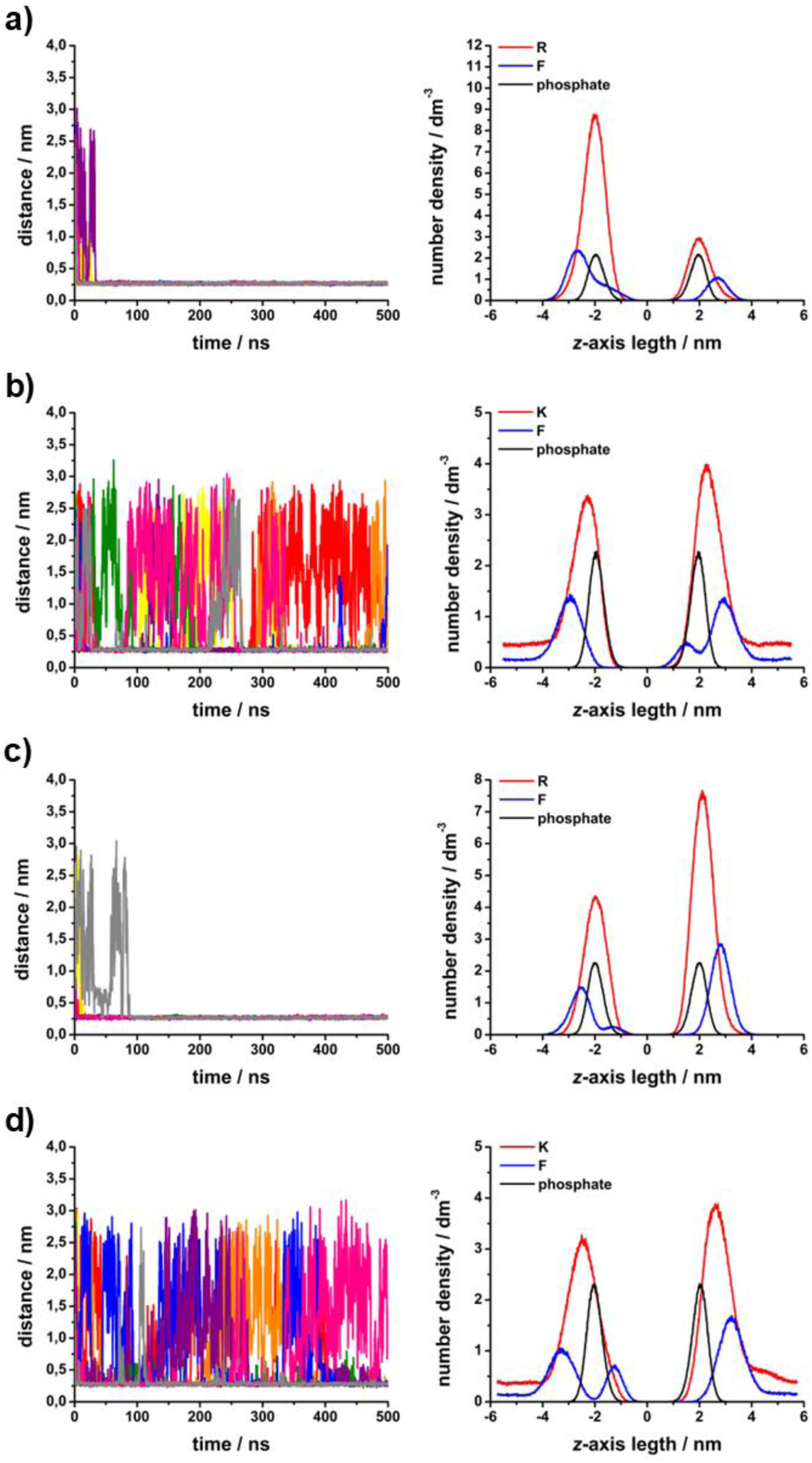
Left: minimum distance of any peptide atom to the phosphate atoms of lipids. Right: number density profiles for P-atoms of lipids, and R, K and F residues of peptides. A) PRO + R5F2, b) PRO + K5F2, c) EU + R5F2, d) EU + K5F2. All systems were simulated at 50 °C.

The impact of peptide binding on the polar headgroups was evaluated by counting the hydrogen bonds (HBs) between lipid groups and peptides or water (Table 2). R5F2 established more HBs both with phosphate and C=O groups, both in PRO and EU systems (with the one exception of C=O···R5F2 bonds in EU systems at 30 °C). It also displaced water from the lipid interface and reduced the HBs formed between water and phosphate groups. However, water interactions with C=O groups were significantly less affected, possibly because peptide insertion among the lipid backbone also allows more water to reach it.

**Table 2.**
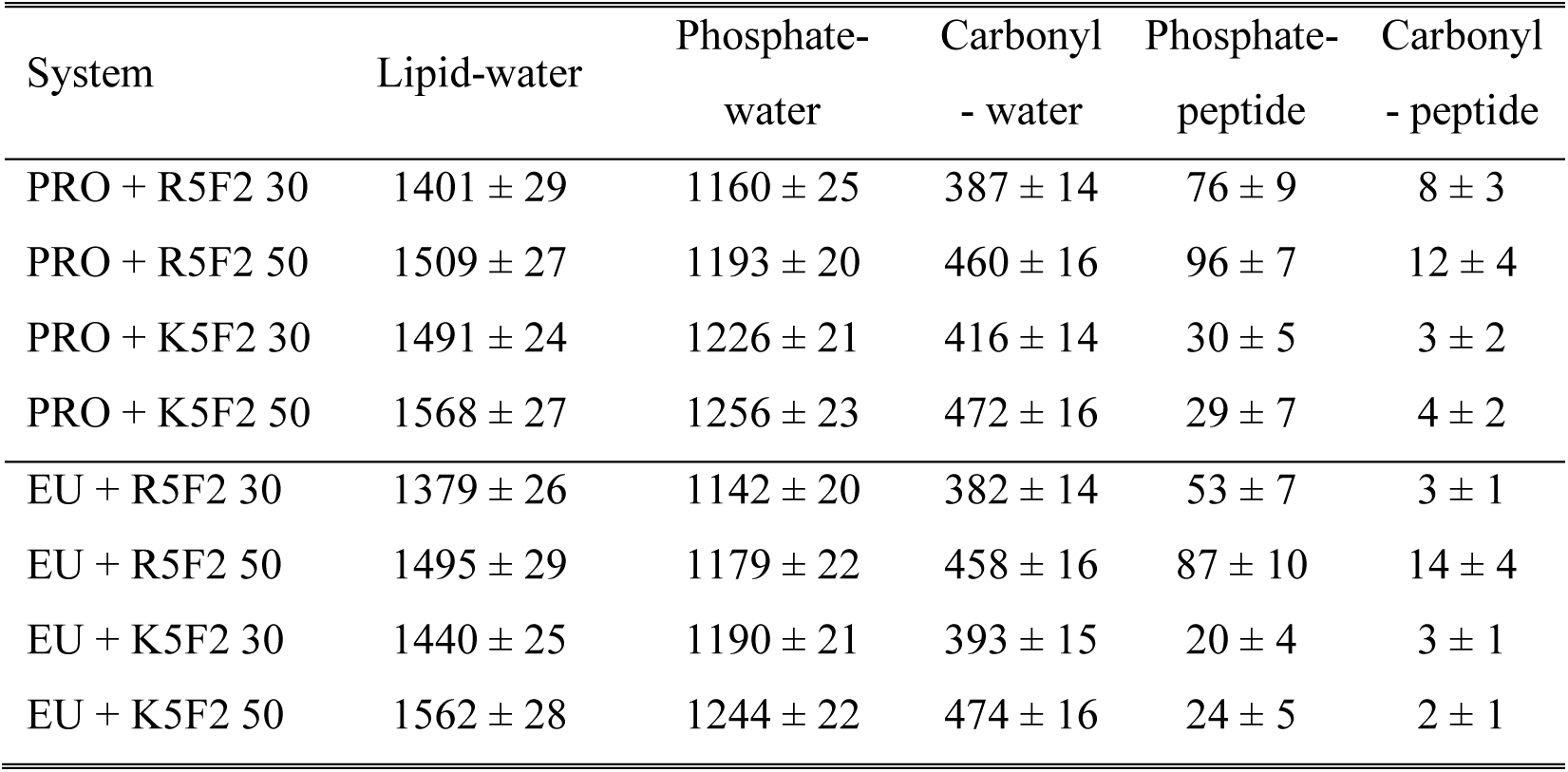
Hydrogen bonds established between select molecules or functional groups for PRO or EU systems in the presence of R5F2 or K5F2, at 30 or 50 °C.

| System | Lipid-water | Phosphate-<br>water | Carbonyl<br>- water | Phosphate-<br>peptide | Carbonyl<br>- peptide |
| --- | --- | --- | --- | --- | --- |
| PRO + R5F2 30 | 1401 ± 29 | 1160 ± 25 | 387 ± 14 | 76 ± 9 | 8 ± 3 |
| PRO + R5F2 50 | 1509 ± 27 | 1193 ± 20 | 460 ± 16 | 96 ± 7 | 12 ± 4 |
| PRO + K5F2 30 | 1491 ± 24 | 1226 ± 21 | 416 ± 14 | 30 ± 5 | 3 ± 2 |
| PRO + K5F2 50 | 1568 ± 27 | 1256 ± 23 | 472 ± 16 | 29 ± 7 | 4 ± 2 |
| EU + R5F2 30 | 1379 ± 26 | 1142 ± 20 | 382 ± 14 | 53 ± 7 | 3 ± 1 |
| EU + R5F2 50 | 1495 ± 29 | 1179 ± 22 | 458 ± 16 | 87 ± 10 | 14 ± 4 |
| EU + K5F2 30 | 1440 ± 25 | 1190 ± 21 | 393 ± 15 | 20 ± 4 | 3 ± 1 |
| EU + K5F2 50 | 1562 ± 28 | 1244 ± 22 | 474 ± 16 | 24 ± 5 | 2 ± 1 |

Thus, the findings from MD display higher tendency of R5F2 to bind and remain bound to the membrane. As was discussed in our previous work [23], this can be attributed to the ability of R residues to form more HBs with the polar groups of the lipids. In contrast to that work, the binding of K residues was enhanced due to the presence of negatively charged lipids in the bilayer. This membrane composition also allowed for peptide penetration into the membrane to be observed in the simulation time. The insertion was expected to be more likely in the fluid phase due to larger gaps in the packing of polar groups. Interestingly, both R5F2 and K5F2 have successfully penetrated the bilayer despite the smaller likelihood of K5F2 binding. R5F2 forms more HBs with phosphate and C=O groups of lipids compared to K5F2, due to the combination of higher likelihood of binding and larger potential of the Gdm^+^ residue for HB formation, in comparison to the NH_3_^+^. The differences between PRO and EU systems were much less pronounced, both in peptide binding and hydration effects, despite the experimental evidence to the contrary. Here it is important to emphasize the limitations of MD, as it only computes a small segment of the bilayer with a few peptide molecules, and ignores other phenomena such as curvature or proton transfer, thus making it harder to observe significant effects.

### 3.4. Microscopy analysis of giant vesicles: stability and partial permeation induced by R5F5

Prompted by the MD simulation results, we explored the effect of R5F2, as the more promising peptide, on model PRO’ membranes. GUVs are ideal cell-sized model membranes due to their ability to mimic the structural and functional properties of biological membranes, allowing for direct microscopy observation of membrane responses, interactions, and processes in a controlled environment [31,70]. We excluded gel-phase membranes from our study due to their typical leakiness observed on GUVs [71].

The vesicles were observed using bright-field and fluorescence microscopy, with the membrane labeled using DilC_18_.Confocal imaging revealed no significant changes in vesicle morphology or population for low concentration (Fig. S6). The GUVs were prepared in sucrose solutions and diluted in glucose, allowing us to monitor changes in vesicle stability and leakiness due to differences in solution densities and refractive indices under bright-field field or phase-contrast observation (Fig. S7).

Contrary to fluorescence observations, which showed no qualitative changes or ruptures, careful inspection of bright-field images revealed leakage, visually recognized by loss of contrast. This was occasionally observed in GUVs subjected to R5F2 at concentrations of 5 μM and higher (see Fig. 6 and Fig. S8). This behavior is reminiscent of the toroidal pore action of antimicrobial peptides [72,73] and is similar to the effects observed with detergents at subsolubilization concentrations [74,75]. Over time, approximately 10% of the vesicles (based on a sample size of over 940 GUVs) exhibited leakage within 30-40 minutes of incubation. In the absence of the peptide, no vesicle permeation was observed during this period. Combined, these observations suggest that R5F5 induces membrane restructuring, decreased packing, and pore formation, that allows exchange of material between the membrane leaflets, and therefore, peptide translocation. These processes also imply effective increase of the water content within the vesicle membrane consistent with the tendency of the peptide to remain bound to the membrane, as well as with detecting more end *gauche* conformers in PRO+R5F2 than in PRO+K5F2. Although using fluorescently labeled peptide analogs to visualize and potentially quantify peptide adsorption and penetration across the membrane would have been be advantageous, the small molecular weight of the peptides suggests that adding a fluorescent tag could significantly impact their performance.

**Fig. 6.**
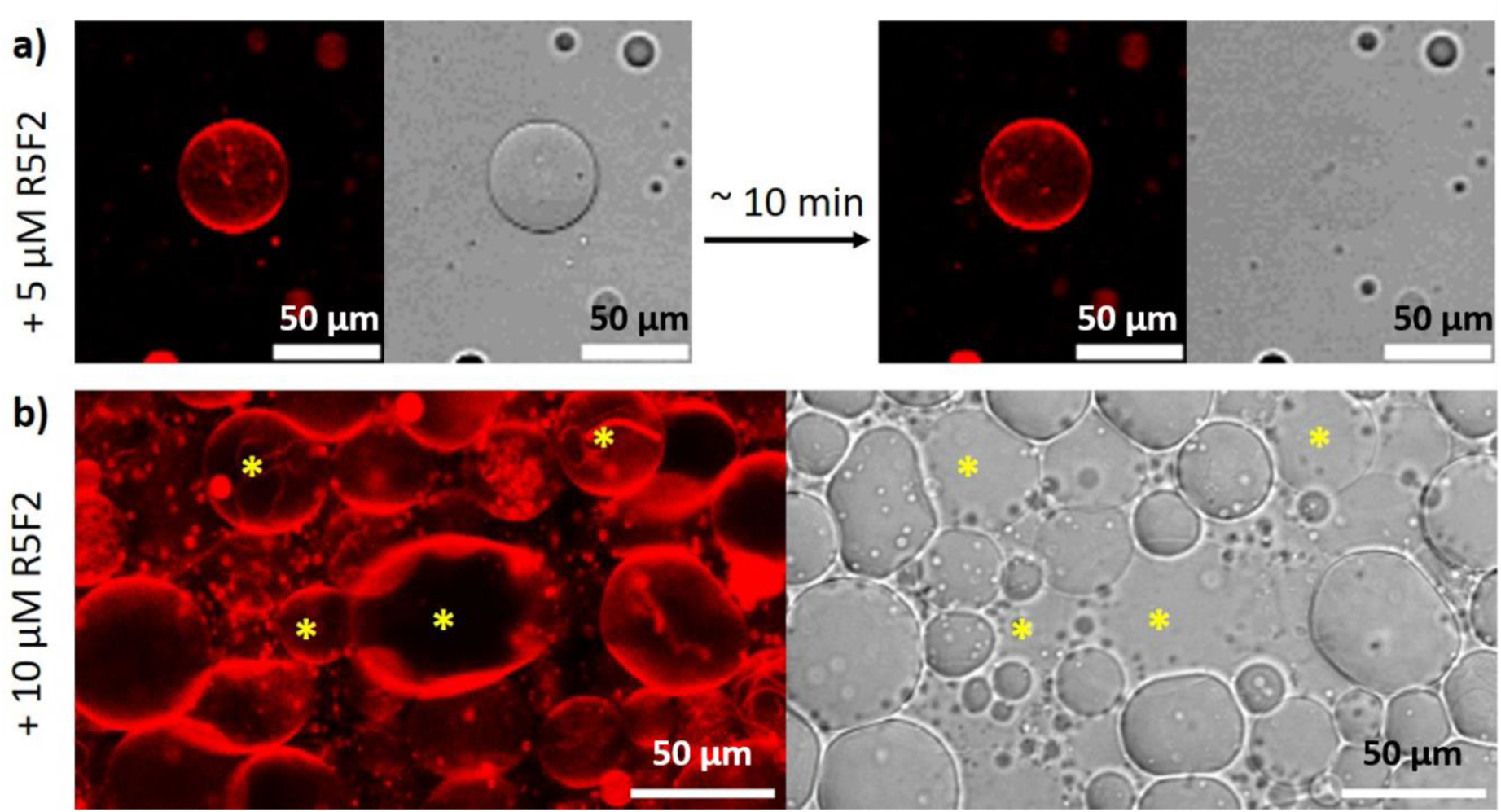
In the presence of R5F2, a fraction of the PRO’ vesicles are susceptible to leakage. In red (left): confocal cross sections; in grey (right): bright field images, displaying changes in contrast caused by sugar asymmetry across the membrane. Images were enhanced to more easily display the variations of contrast. Scale bar 50 µm. a) Single-vesicle observation over time. Snapshots displaying the loss of contrast and thus leakage of the GUV after the addition of 5 µM R5F2. b) Bulk observation over a population of GUV displaying multiple leaky GUVs several minutes after the addition of 10 µM R5F2.

## 4. Discussion

The examination of all obtained results can be summarized in the following main findings (going from the top of the lipid bilayer to below):

i) When approaching polar moieties of PRO system, i.e. dominantly choline and especially of phosphate groups, R5F2 simultaneously decreases the transition dipole moment of ν_s_PO_2_^−^ band and increases that of ν_as_CN. This significant charge redistribution probably emerges due to the orientation of R5F2 against lipid polar moieties and overall penetration depth. This phenomenon does not resemble the response of K5F2 since in PRO+K5F2, as well as in EU+K5F2, where one cannot see so dramatic changes in the relative intensities; i.e. although the intensity of ν_s_PO_2_^−^ band is affected by the presence of K5F2, but not as the reduction and high-frequency displacement of the νCOP band (ν_as_CN band appears to be not very sensitive to this). Presumably, upon the interaction of K5F2 a shrinkening of νCOP moieties occurs, likely due to reduced hydration in the presence of K5F2. Since EU system in both the absence and presence of R5F2 produces qualitatively similar responses of these bands, R5F2 do not directly interact with these lipid parts. However, in the EU+K5F2 system the broadening of νCOP band on its high-frequency side, although not merged as in PRO+K5F2, may be related with a decreased hydration of COP moiety in the presence of K5F2. The presumption of COP moiety as a hotspot for interaction with peptides, accompanied by a significant change in the hydration in the presence of K5F2, goes along with relatively poor changes in the response of ν_as_PO ^−^ band; in PRO+R5F2/K5F2 systems the maximum remains at the same place all the time, whereas in EU+R5F2/K5F2 it slightly displaces to lower frequencies, suggesting that gel-to-fluid phase transition of EU systems in the presence of peptides is very likely accompanied with increased hydration in comparison with PRO systems. The MD simulations support this assertion, with the higher positive difference (gain) in HBs between gel and fluid in EU systems, but they did not reproduce the dehydration caused by K5F2.

ii) As far as polar-apolar interface is concerned (νC=O band), the reduced hydration of PRO system in the presence of R5F2, unlike that of K5F2, suggests the possibility that R5F2 interacts directly with carbonyl moiety and thus prevents the approach of water molecules allowed by K5F2 (the difference between pure PRO response and that of PRO+K5F2 is negligible). On the other hand, the similarity in responses of EU ± R5F2/K5F2 suggest the absence of any dramatic changes caused by the presence of peptides. MD simulations confirmed more HB formation between C=O and R5F2 compared to K5F2, except in the case of EU systems at 30 °C. Before going deeper into the non-polar part of lipid bilayers, we would like to draw attention to the ζ-potential values. For PRO and EU systems they are −6.5 ± 0.4 mV and −16.0 ± 0.3 mV, due to the presence of negatively charged lipids. The presence of R5F2 leads to ζ-potential increase for about 2 mV, i.e. −4.5 ± 0.9 mV in PRO+R5F2 and −13.6 ± 0.5 mV in EU+R5F2, suggesting charge screening by the adsorption of the peptide. Importantly, in the presence of K5F2, the ζ-potential of the PRO system decreases to −10.0 ± 0.8 mV, whereas the analogous EU system significantly increases and effectively approaches 0, suggesting that, in addition to charge screening from adsorption, there might be proton transfer between COO^−^ group of DPPS and −NH_3_^+^ of K5F2 [15,23]. This phenomenon might also contribute to the observed high-frequency displacement of inherently broad and towards lower frequencies skewed νC=O_(non-)HB_ band generated by lipid molecules that ought to overlap νC=O band generated by COOH group expected at about 1710 cm^−1^ [76]. The decrease in ζ-potential of PRO+K5F2 may be associated with the reduction in the thickness of the hydration shell around the liposome. Nevertheless, at this point we have to refer to the results obtained from temperature-dependent UV-Vis and DSC responses: in general, it is fair to say that pretransition is barely detectable from DSC curves, except for the systems that include K5F2 (see Fig. 3 and Table 1). Moreover, UV-Vis data suggest that only K5F2 in both systems enhances the occurrence/magnitude of ripples at the bilayer surface. In this light, it seems reasonable to assume that K5F2 remains closer to the surface of the bilayer and induces substantial charge redistribution within terminal polar moieties of lipid molecules, whereas R5F2 may penetrate a little bit deeper in the bilayer. Though MD simulations have shown that both peptides may penetrate the membrane to reach the hydrophobic segment, considering the binding of R5F2 is more likely and more stable, it is reasonable to assume it will have more prominent effects deeper in the membrane. The microscopy observations showing partial GUV leakage induced by R5F2 (Fig. 6) might imply full insertion of the peptide, leading to membrane restructuring and pore formation. This process effectively increases the water content and disorganizes the lipids consistent with our observations of decresed packing.

iii) The γCH_2_ bands, mirroring the lateral interactions between hydrocarbon chains, clearly imply that the presence of peptides alters the packing pattern. Putting this differently, the adsorption of peptides impacts the conformation of glycerol moiety and the mutual position and orientation of hydrocarbon chains [60,77,78]. This seems to be particularly pronounced in PRO+R5F2 since the latter modulates the packing pattern inherent for pure PRO lipid bilayers, i.e. from orthorhombic in the gel phase to hexagonal. The difference in the packing pattern of PRO and EU systems is also distinguished from their thermotropic data, especially obtained from DSC (see Table 1) since PRO systems generally provide *T*_m (1/2)_ smaller for about 1 °C than the analogous quantity in EU systems [67,79]. Ultimately, crucial differences in the action of these two peptides onto lipid bilayer emerge from the features of ωCH_2_ bands. Owing to the differences in the intensities of the bands associated with double *gauche*, end *gauche* and kink conformers, it can be assumed that the interaction of polar lipid parts with peptides reflects on the number and distribution of these conformers across the hydrophobic part. In this context, a greater number of double *gauche* conformers in PRO+K5F2 (owing to the stronger band associated with them), in contrast to more end *gauche* conformers in PRO+R5F2 (Fig. 4g), might imply that K5F2 induces smaller number of larger defects in PRO system than R5F2, but in the presence of the latter the inversion appears, i.e. a larger number of smaller defects within non-polar bilayer part. Knowing that these peptides do not disrupt the vesicles overall (Fig. 6 and Figs. S6-S8), these defects may be considered as potential vacancies [28] occasionally leading to pore formation. The analogous phenomenon in EU systems is not so pronounced. The potential contribution of proton transfer between DPPS and K5F2 [15] on the type and distributions of *gauche* conformers [28], as well as on the detection of ripple phase [80] cannot be excluded but is out of the scope of the present study.

One more phenomenon ought to be more deeply elaborated. According to Cevc, ripple phase in DPPC lipid (multi)bilayers is a result of a longitudinal displacement of lipid molecules owing to a different hydration pattern in certain temperature range [37]. Besides more frequent adsorption/desorption of K5F2 from both PRO and EU bilayer surface, there is a possibility that K5F2 induces hydration changes in the interfacial water layer that reinforce this longitudinal displacement of lipid molecules, increasing their protrusion beyond the surface. Moreover, the lack of signals originated from the vibrations of glycerol moiety of DPPG lipids in PRO systems in the presence of R5F2 (Fig. 4c) might go along with the suppression of longitudinal displacement of the lipids in the presence of R5F2. In this light, one can conclude that the adsorption of peptides onto PRO and EU lipid bilayers is dominantly driven by the charge distribution of peptides, implying that the net charge of lipid bilayers is of secondary importance. The increased longitudinal displacement in the presence of K5F2, but not in R5F2, should be mirrored in the occurrence of different *gauche* conformers (Fig. 4g), especially in intrinsically less hydrated gel phase of lipid bilayers than fluid phase [56,57,81]. Assuming that the *gauche* conformers can be taken as defects, these results can serve as a platform for deciphering how the change in the hydration pattern and sliding the lipids along the membrane normal affects the type and distribution of vacancies that might serve as a pathway for peptide translocation.

## 5. Conclusions

Both R5F2 and K5F2 show a propensity for binding to the membrane, though R5F2 has a higher affinity and greater ability to interact with the polar groups and form hydrogen bonds. Depending on the composition of the lipid membrane (PRO or EU) the peptides exerted a different effect on the membrane properties; in particular, their *T*_m_ values (DSC) are modulated in a way that in PRO/EU systems the softening/rigidification of lipid bilayers is detected.

Turbidity-based data (UV-Vis) that detect ripple phase (*T*_p_ determination) only when K5F2 are present in the systems might be related to a considerable impact of K5F2 on the interfacial water layer on both PRO and EU systems. Though both peptides possess membrane-penetrating potential, R5F2 is likely situated deeper in the membrane, while K5F2 retains positions at the surface, and possibly longitudinally displaces lipids. Most importantly, the detection of peptide-dependent *gauche* conformers for PRO systems that serve as models for bacterial membranes, along with the maintenance of their integrity, implies that the number and type of *gauche* conformers may be viewed as precursors of defects that will ultimately, in a concerted manner, enable peptide translocation across lipid bilayers.

## Supporting Information

Supporting Information associated with this paper can be found in the online version at http://.

## Acknowledgment

This paper was supported by Croatian Science Foundation, Project No. UIP-2020-02-7669 and by a bilateral project “Myelin protein impact on membrane phase state, morphology and structure (ImProMem)” financed by Croatian Ministry of Science and Education (MSE) and Deutscher Akademischer Austauschdienst (DAAD). L. P. and D. B. sincerely thank the members of Laboratory for biocolloids and surface chemistry (Division for Physical Chemistry, Ruđer Bošković Institute) for who enabled access to the Zetasizer Nano ZS (Malvern). L. P., A. J. and D. B. thank Mrs. Milica Perc (Ruđer Bošković Institute) for the assistance in peptides purification and Dr. Amela Hozić and Dr. Mario Cindrić for HRMS measurements and analyses (Ruđer Bošković Institute). B. P. thanks Dr Dražen Petrov (BOKU University, Vienna, Austria) for helpful comments regarding the molecular dynamics setup.

