## Supporting Information for "Peptide-induced hydration of lipid bilayers modulates packing pattern and conformations of hydrocarbon chains – a potential pathway for peptide translocation?"

**S1. R5F2/K5F2: Spectral characterization** p2

**S2. DLS and ζ-potential data: PRO ± R5F2/K5F2 and EU ± R5F2/K5F2** p4

**S3. DSC curves of PRO ± R5F2/K5F2 and EU ± R5F2/K5F2 as MLVs** p5

**S4. FTIR spectra of R5F2 and K5F2** p6

**S5. Additional molecular dynamics data** p7

**S6. Additional microscopy data of GUVs** p10

**References** p13

### S1. R5F2/K5F2: Spectral characterization

The synthesis of R5F2 and K5F2 on a solid phase is thoroughly described in our previous paper [1]. Here, only the data obtained from NMR spectra are listed in the text below, proving their structure.

#### *S1.1 Characterization of K5F2 and R5F2 peptides*

*L-lysyl-L-lysyl-L-lysyl-L-lysyl-L-lysyl-L-phenylalanyl-L-phenylalanine (KKKKKFF,* ***K5F2)***

**$^1$H NMR** (600 MHz, DMSO-*d*$_6$): $\delta$ = 1.11 - 1.18 (m, 2 H, Lys$^5$-γ), 1.18 - 1.22 (m, 2 H, Lys$^4$-γ), 1.24 - 1.28 (m, 2 H, Lys$^3$-γ), 1.28 - 1.33 (m, 2 H, Lys$^2$-γ), 1.33 - 1.38 (m, 2 H, Lys$^1$-γ), 1.43 - 1.49 (m, 3 H, Lys$^{3-5}$-β''), 1.49 - 1.55 (m, 12 H, Lys$^{1-5}$-γ, Lys$^2$-β'', Lys$^5$-β'), 1.56 - 1.61 (m, 1 H, Lys$^2$-β'), 1.60 - 1.66 (m, 2 H, Lys$^{2,3}$-β'), 1.66 - 1.74 (m, 2 H, Lys$^1$-β), 2.65 - 2.80 (m, 11 H, Lys$^{1-5}$-ε, Phe$^6$-β''), 2.92 (dd, *J* = 14.1, 8.6 Hz, 1 H, Phe$^7$-β''), 2.99 (dd, *J* = 13.8, 4.2 Hz, 1 H, Phe$^6$-β'), 3.07 (dd, *J* = 14.1, 5.3 Hz, 1 H, Phe$^7$-β'), 3.81 (br dd, *J* = 11.7, 5.9 Hz, 1 H, Lys$^1$-α), 4.15 (td, *J* = 7.8, 5.0 Hz, 1 H, Lys$^5$-α), 4.17 - 4.20 (m, 1 H, Lys$^4$-α), 4.20 - 4.25 (m, 1 H, Lys$^3$-α), 4.29 (td, *J* = 8.2, 5.3 Hz, 1 H, Lys$^2$-α), 4.45 (td, *J* = 8.3, 5.3 Hz, 1 H, Phe$^7$-α), 4.53 (td, *J* = 8.8, 4.4 Hz, 1 H, Phe$^6$-α), 7.16 - 7.18 (m, 1 H, Phe$^6$-ζ), 7.18 - 7.20 (m, 1 H, Phe$^7$-ζ), 7.19 - 7.23 (m, *J* = 2.2 Hz, 4 H, Phe$^6$-δ, Phe$^6$-ε), 7.23 (d, *J* = 7.7 Hz, 2 H, Phe$^7$-δ), 7.27 (t, J = 7.3 Hz, 2 H, Phe$^7$-ε), 7.81 - 7.91 (m, 10 H, Lys$^{1-5}$-ε-$NH_2$), 7.88 - 7.91 (m, 1 H, Phe$^6$-α-NH), 7.96 (br d, *J* = 8.1 Hz, 1 H, Lys$^5$-α-NH), 7.96 (br d, *J* = 7.3 Hz, 1 H, Lys$^4$-α-NH), 8.15 (br d, *J* = 7.7 Hz, 1 H, Lys$^3$-α-NH), 8.22 (br d, *J* = 3.7 Hz, 2 H, Lys$^1$-α-NH), 8.36 (d, *J* = 8.1 Hz, 1 H, Phe$^7$-α-NH), 8.59 (br d, J = 7.3 Hz, 1 H, Lys$^2$-α-NH) ppm.

**$^{13}$C NMR** (151 MHz, DMSO-*d*$_6$): $\delta$ = 21.0 (Lys$^1$-γ-$CH_2$), 22.1 (Lys$^5$-γ-$CH_2$), 22.2 (Lys$^{2-4}$-γ-$CH_2$), 26.3 (Lys$^1$-δ-$CH_2$), 26.6 (Lys$^{2-5}$-δ-$CH_2$), 30.4 (Lys$^1$-β-$CH_2$), 31.4 (Lys$^{2-5}$-β-$CH_2$), 36.7 (Phe$^7$-β-$CH_2$), 37.6 (Phe$^6$-β-$CH_2$), 38.4 (Lys$^1$-ε-$CH_2$), 38.6 (Lys$^{2-5}$-ε-$CH_2$), 51.7 (Lys$^1$-α-CH), 52.2 (Lys$^3$-α-CH), 52.3 (Lys$^{4,5}$-α-CH), 52.5 (Lys$^2$-α-CH 53), 53.3 (Phe$^6$-α-CH), 53.4 (Phe$^7$-α-CH), 126.2 (Phe$^6$-ζ-CH), 126.4 (Phe$^7$-ζ-CH), 127.9 (Phe$^6$-ε-CH), 128.2 (Phe$^7$-ε-CH), 129.1 (Phe$^7$-δ-CH), 129.2 (Phe$^6$-δ-CH), 137.3 (Phe$^7$-γ-C), 137.4 (Phe$^6$-γ-C), 168.4 (Lys$^1$-CO), 170.9 (Lys$^4$-CO), 171.0 (Phe$^6$-CO), 171.1 (Lys$^5$-CO), 171.2 (Lys$^2$-CO), 171.4 (Lys$^3$-CO), 172.6 (Phe$^7$-COOH) ppm.

**$C_{48}H_{80}N_{12}O_8$ *M*r = 952.62, [M + H]$^+$ . HRMS: *m*/*z* 953.6298 [M + H]$^+$ (calc. 953.6300).**

*L-arginyl-L-arginyl-L-arginyl-L-arginyl-L-arginyl-L-phenylalanyl-L-phenylalanine (RRRRRFF, **R5F2**)*

**$^1$H NMR** (600 MHz, DMSO-$d_6$): $\delta$ = 1.36 (br s, 2 H, Arg$^5$-γ), 1.40 - 1.46 (m, 2 H, Arg$^4$-γ), 1.46 - 1.50 (m, 2 H, Arg$^3$-γ), 1.50 - 1.56 (m, 6 H, Arg$^{1,2}$-γ, Arg$^5$-β), 1.58 (br s, 2 H, Arg$^4$-β), 1.62 (br s, 2 H, Arg$^3$-β), 1.69 (br s, 4 H, Arg$^{1,2}$-β), 2.75 (br dd, J = 13.9, 9.2 Hz, 1 H, Phe$^6$-β''), 2.92 (dd, $J$ = 13.9, 8.8 Hz, 1 H, Phe$^7$-β''), 3.00 (dd, $J$ = 13.6, 4.0 Hz, 1 H, Phe$^6$-β'), 3.02 - 3.05 (m, 4 H, Arg$^{4,5}$-γ), 3.05 - 3.07 (m, 3 H, Arg$^3$-γ, Phe$^7$-β'), 3.09 (br d, $J$ = 6.6 Hz, 2 H, Arg$^2$-δ), 3.11 (br s, 2 H, Arg$^1$-δ), 3.84 (dt, $J$ = 7.0, 6.0 Hz, 1 H, Arg$^1$-α), 4.16 - 4.20 (m, 1 H, Arg$^5$-α), 4.22 (br d, J = 7.0 Hz, 1 H, Arg$^4$-α), 4.23 - 4.26 (m, 1 H, Arg$^3$-α), 4.33 (dt, $J$ = 8.0, 6.0 Hz, 1 H, Arg$^2$-α), 4.44 (td, $J$ = 8.2, 5.7 Hz, 1 H, Phe$^7$-α), 4.54 (td, $J$ = 8.6, 4.8 Hz, 1 H, Phe$^6$-α), 7.14 - 7.18 (m, 1 H, Phe$^6$-ζ), 7.18 - 7.21 (m, 5 H, Phe$^6$-δ,ε, Phe$^7$-ζ), 7.22 (br d, $J$ = 8.4 Hz, 2 H, Phe$^7$-δ), 7.26 (t, $J$ = 7.0 Hz, 2 H, Phe$^7$-ε), 7.29 - 7.53 (m, 15 H, Arg$^{1\text{-}5}$-ε-NH,$NH_2$), 7.65 (br t, $J$ = 5.0 Hz, 1 H, Arg$^5$-γ-NH), 7.70 (br t, $J$ = 5.1 Hz, 1 H, Arg$^4$-γ-NH), 7.79 (br t, $J$ = 5.5 Hz, 1 H, Arg$^3$-γ-NH), 7.85 (br t, $J$ = 5.9 Hz, 1 H, Arg$^1$-γ-NH), 7.86 (t, $J$ = 6.2 Hz, 1 H, Arg$^2$-γ-NH), 7.95 (br d, $J$ = 8.1 Hz, 1 H, Phe$^6$-α-NH), 8.02 (d, $J$ = 7.7 Hz, 1 H, Arg$^5$-α-NH), 8.03 (d, $J$ = 7.0 Hz, 1 H, Arg$^4$-α-NH), 8.19 (br d, $J$ = 7.7 Hz, 1 H, Arg$^3$-α-NH), 8.22 (br d, $J$ = 3.7 Hz, 2 H, Arg$^1$-α-$NH_2$), 8.44 (br d, $J$ = 7.7 Hz, 1 H, Phe$^7$-α-NH), 8.60 (br d, $J$ = 7.3 Hz, 1 H, Arg$^2$-α-NH) ppm.

**$^{13}$C NMR** (151 MHz, DMSO-$d_6$): $\delta$ = 24.0 (Arg$^1$-γ-$CH_2$), 24.8 (Arg$^5$-γ-$CH_2$), 24.9 (Arg$^2$-γ-$CH_2$), 25.0 (Arg$^{3,4}$-γ-$CH_2$), 28.4 (Arg$^1$-β-$CH_2$), 29.1 (Arg$^5$-β-$CH_2$), 29.2 (Arg$^{3,4}$-β-$CH_2$), 29.3 (Arg$^2$-β-$CH_2$), 36.7 (Phe$^7$-β-$CH_2$), 37.6 (Phe$^6$-β-$CH_2$), 40.1 (Arg$^1$-δ-$CH_2$), 40.4 (Arg$^{2\text{-}5}$-δ-$CH_2$), 51.7 (Arg$^1$-α-CH), 52.1 (Arg$^{4,5}$-α-CH), 52.2 (Arg$^3$-α-CH), 52.4 (Arg$^2$-α-CH), 53.3 (Phe$^6$-α-CH), 53.6 (Phe$^7$-α-CH), 126.2 (Phe$^6$-ζ-CH), 126.4 (Phe$^7$-ζ-CH), 127.9 (Phe$^6$-ε-CH), 128.1 (Phe$^7$-ε-CH), 129.1 (Phe$^6$-δ-CH), 129.2 (Phe$^7$-δ-CH), 137.3 (Phe$^{6,7}$-γ-CH), 156.8 (Arg$^{1\text{-}5}$-δ-NH), 168.4 (Arg$^1$-CO), 170.9 (Arg$^{2,4}$-CO), 171.0 (Arg$^5$-CO), 171.1 (Phe$^6$-CO), 171.3 (Arg$^3$-CO), 172.6 (Phe$^7$-COOH) ppm.

**$C_{48}H_{80}N_{22}O_8$ *M*r = 1093.66, $[M + H]^+$. HRMS: *m*/*z* 1093.6608 $[M + H]^+$ (calc. 1093.608).**

**S2. DLS and ζ-potential data: PRO ± R5F2/K5F2 and EU ± R5F2/K5F2**

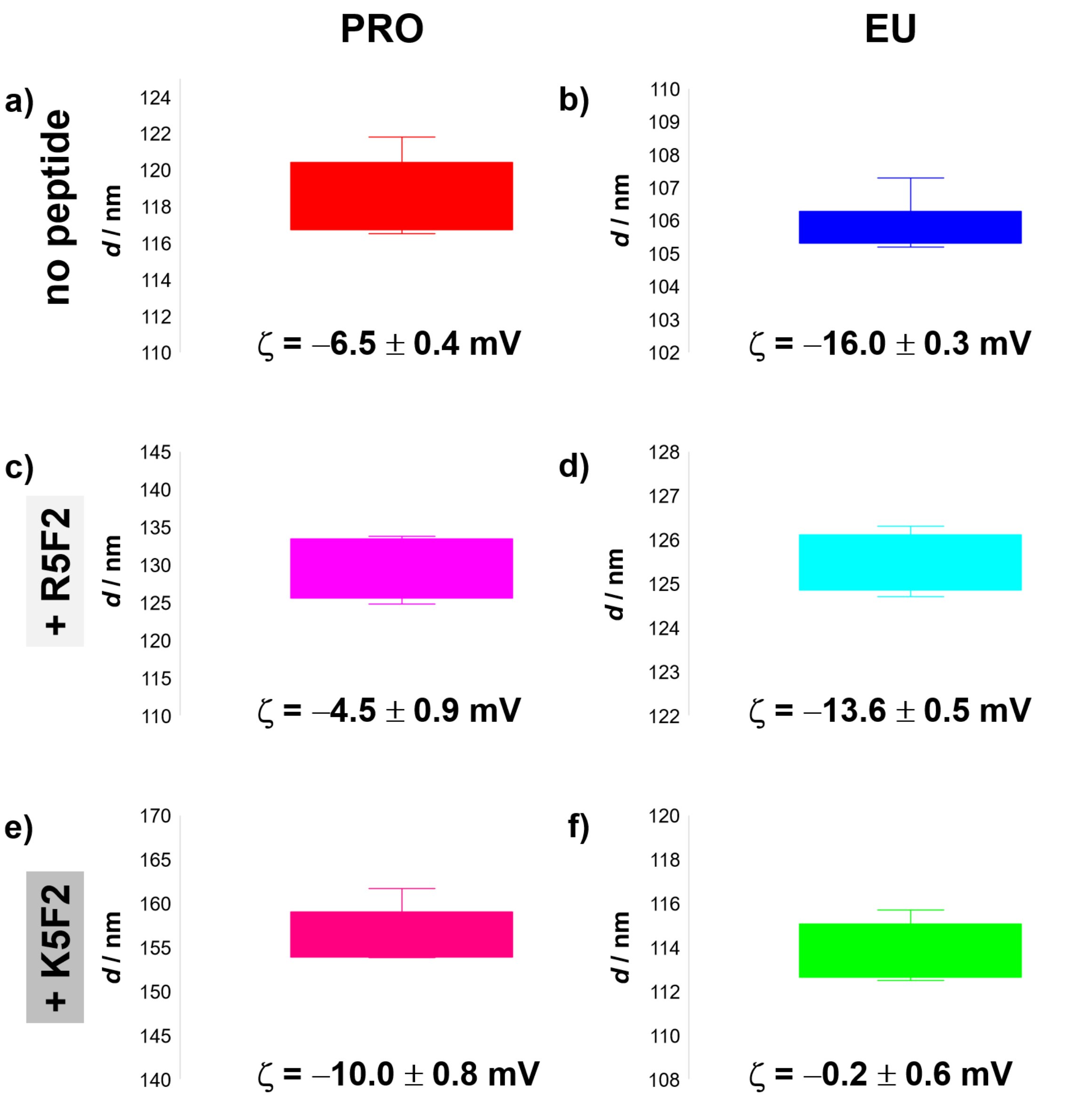


Fig. S1. DLS and ζ-potential data of: a) PRO; b) EU; c) PRO+R5F2; d) EU+R5F2; e) PRO+K5F2; f) EU+K5F2 obtained at 25 °C.

**S3. DSC curves of PRO ± R5F2/K5F2 and EU ± R5F2/K5F2 as MLVs**

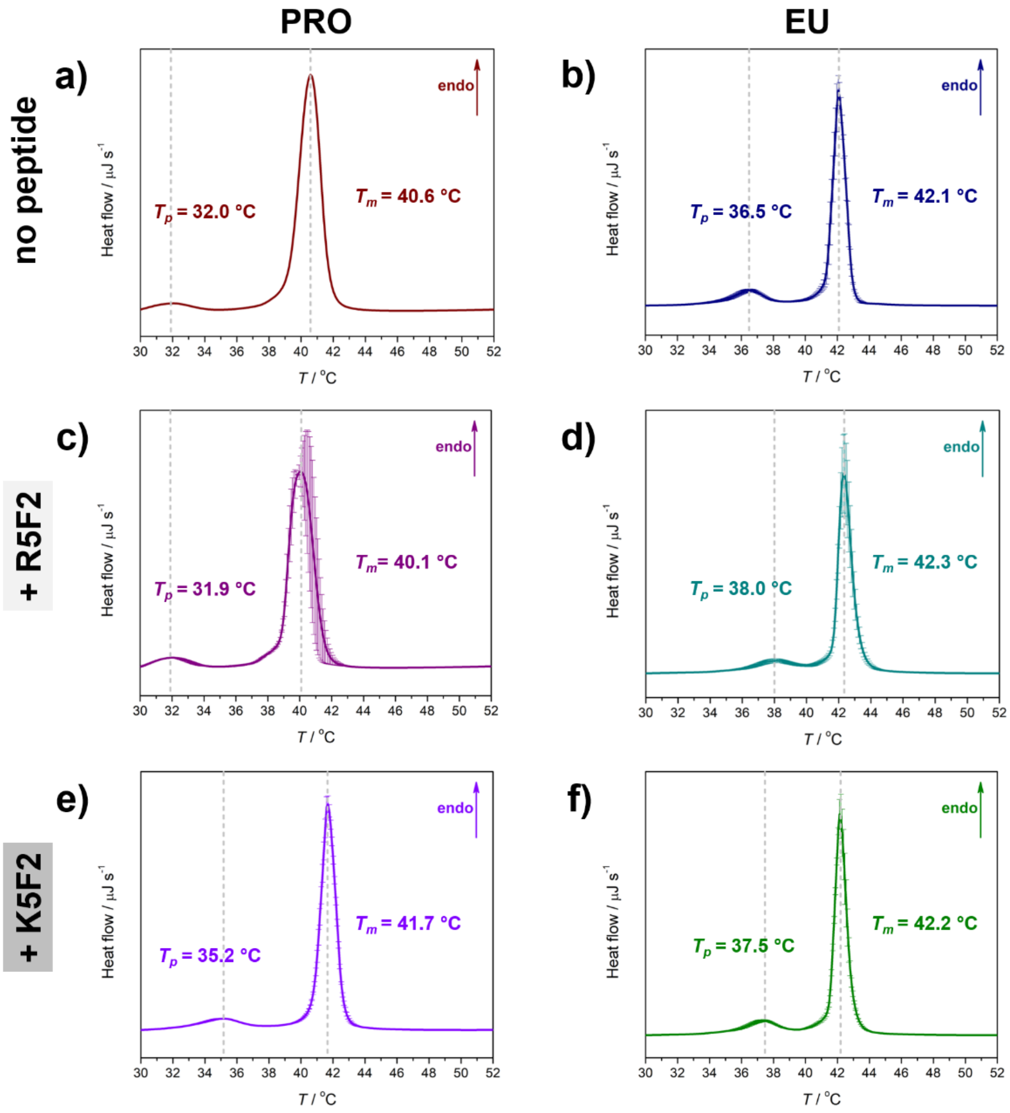


Fig. S2. DSC curves of: a) PRO; b) EU; c) PRO+R5F2; d) EU+R5F2; e) PRO+K5F2; f) EU+K5F2 obtained in the 2nd heating run in the form of MLVs.

**S4. FTIR spectra of R5F2 and K5F2**

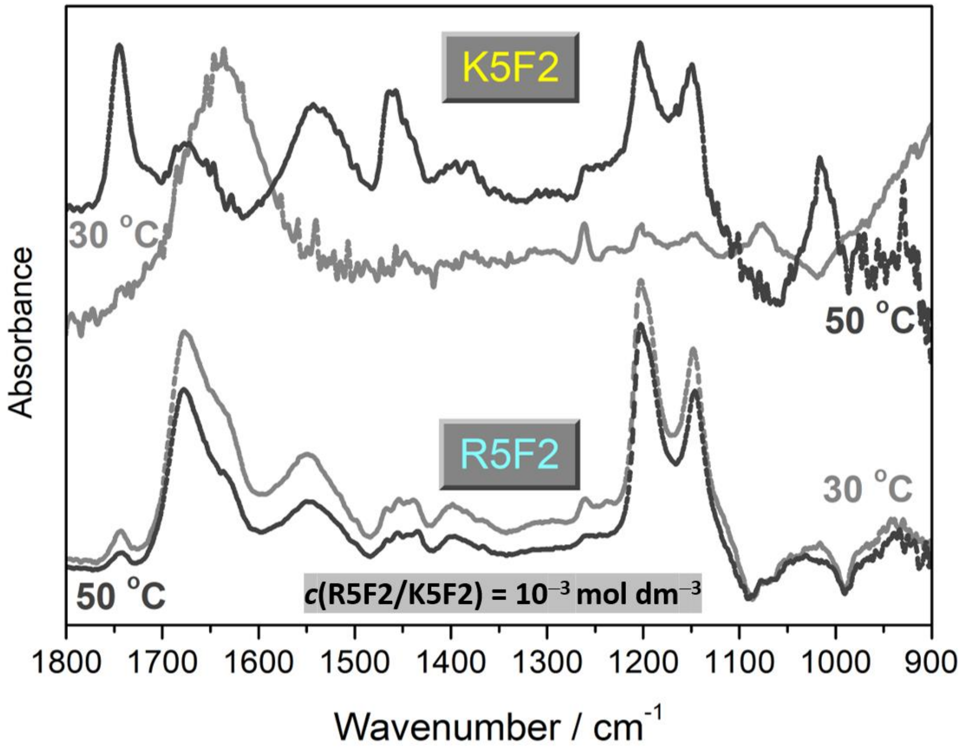


Fig. S3. FTIR spectra of R5F2/K5F2 after PB subtraction at 30 °C and 50 °C (*c*(R5F2/K5F2) = 0.001 mol $dm^{-3}$).

### S5. Additional molecular dynamics data

Area per lipid (APL) is determined as the *xy* area of the simulation box divided by the number of lipids in a single leaflet. It is an indicator of lipid packing, with higher values denoting the fluid phase. As seen in Table APL, the values are similar to the experimentally determined APL of DPPC (0.46 ± 0.02 $nm^2$ in gel and 0.62 ± 0.02 $nm^2$ in fluid) [2,3]. The increase in APL is accompanied by the decrease in membrane thickness (MT, Table S1). MT is reported as the average distance of P-atoms in opposing leaflets, taken from the phosphorus number density distribution. The values are higher than previously reported for gel phase [4,5], likely due to the formation of ripples. Rippling was observed in the simulations at 30 °C, which is lower than the experimental $T_p$, but was noted feature of DPPC simulations [6]. The deuterium order parameters (Fig. S4), showing the level of organization of acyl chains, demonstrate high structuring (large -$S_{CD}$) at 30 °C, and high disorder (low -$S_{CD}$) at 50 °C, confirming the phase difference.

Table S1. Area per lipid (APL) and membrane thickness (MT) of DPPC + 10% DPPG (PRO) or DPPS (EU) systems in the presence of K5F2 or R5F2 peptides, at 30 °C or 50 °C.

| System | APL / $nm^2$ | MT / nm |
|---|---|---|
| PRO + R5F2 30 | 0.482 ± 0.004 | 5.163 ± 0.099 |
| PRO + R5F2 50 | 0.619 ± 0.009 | 3.896 ± 0.126 |
| PRO + K5F2 30 | 0.501 ± 0.005 | 5.333 ± 0.110 |
| PRO + K5F2 50 | 0.617 ± 0.010 | 3.895 ± 0.013 |
| EU + R5F2 30 | 0.488 ± 0.005 | 5.102 ± 0.066 |
| EU + R5F2 50 | 0.608 ± 0.011 | 4.033 ± 0.014 |
| EU + K5F2 30 | 0.497 ± 0.004 | 4.580 ± 0.147 |
| EU + K5F2 50 | 0.609 ± 0.010 | 4.003 ± 0.069 |

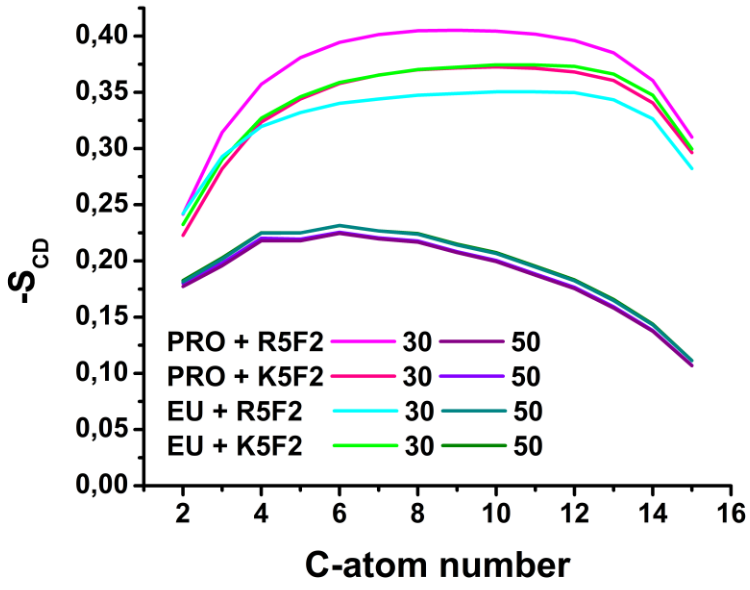


Fig. S4. Deuterium order parameters of acyl chains for the select systems at 30 °C or 50 °C.

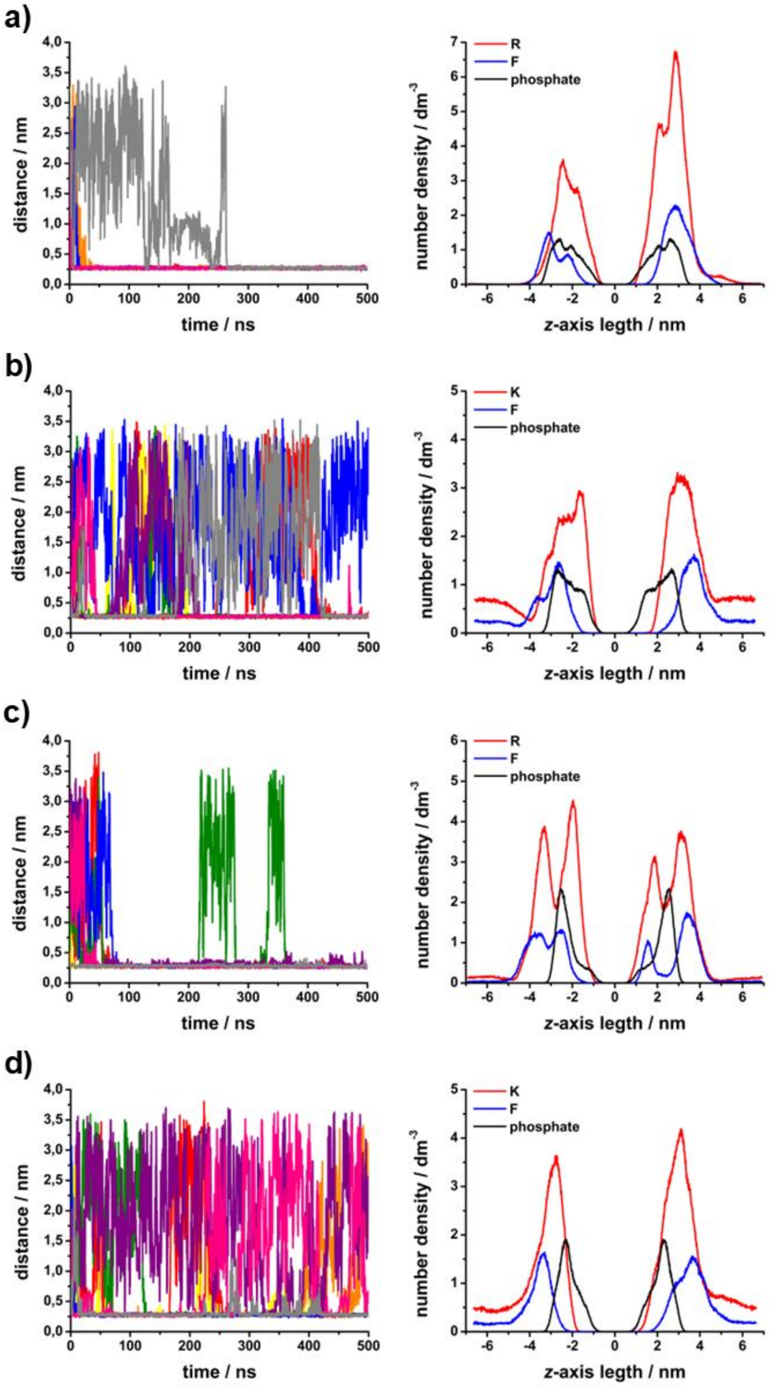


Fig. S5. Left: minimum distance of any peptide atom to the phosphate atoms of lipids. Right: number density profiles for P-atoms of lipids, and R, K and F residues of peptides. A) PRO + R5F2, b) PRO + K5F2, c) EU + R5F2, d) EU + K5F2. All systems were simulated at 30 °C.

### S6. Additional microscopy data of GUVs

The images of GUVs representing PRO’ and EU’ in the absence/presence of R5F2/K5F2 with smaller concentrations of peptides (1 µM) are displayed in Fig. S6. A closer inspection of PRO’ GUVs following the addition of the peptide-free buffer did not show a loss of contrast (Fig. S7).

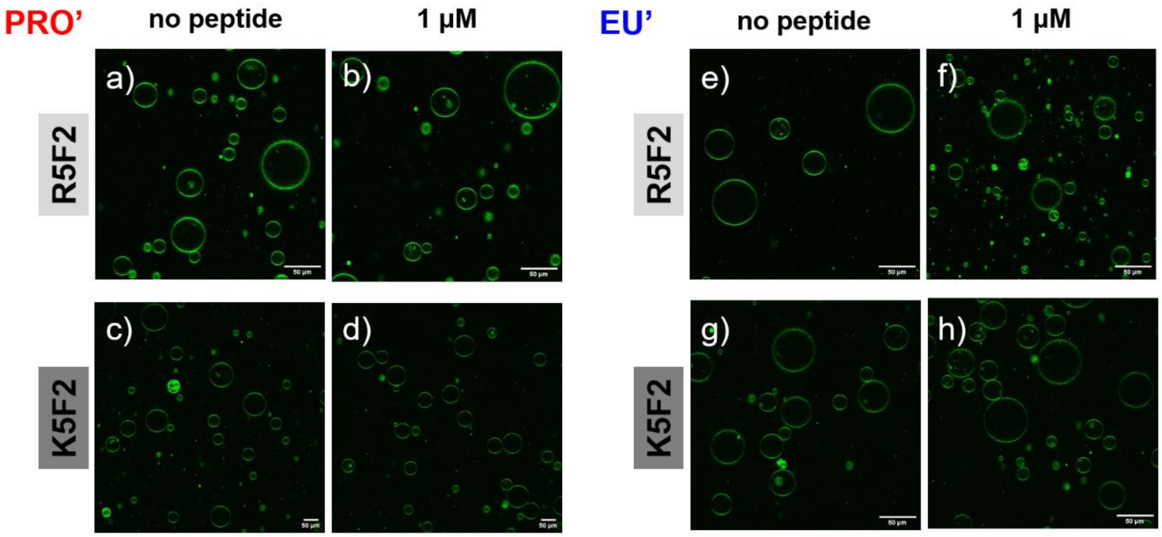


Fig. S6. Confocal images of: a, c) PRO’ GUVs; b, d) PRO’ GUVs in the presence of 1 µM R5F2 and K5F2; e, g) EU’ GUVs; f, h) EU’ GUVs in the presence of 1 µM R5F2 and K5F2.

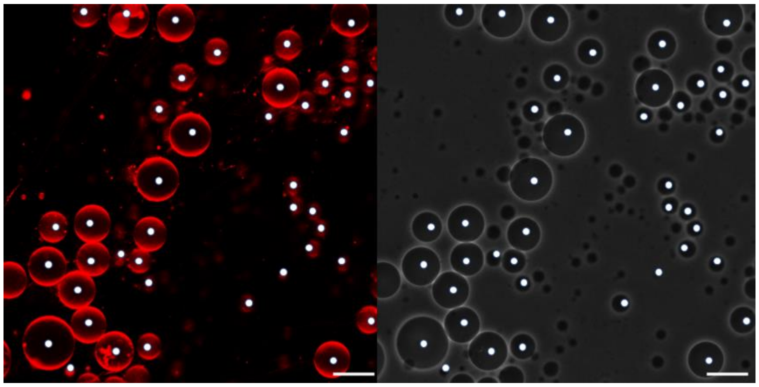

Fig. S7. Images of PRO GUVs recorded on a Leica SP8 confocal microscope. In red (left): confocal microscopy; in grey (right): phase contrast, displaying phase contrast caused by sugar asymmetry. Contrast was enhanced in the images to more easily display the variations of phase contrast. Scale bar 50 µm. The GUVs marked with a white circle in the fluorescence channel show that they are also visible in the phase contrast image, meaning that they have not become leaky >10 minutes after the addition of the control buffer.

While the addition of 1 µM R5F2 did not display PRO' GUVs leaking, multiple instances of leakage could be observed after adding 5 µM (Fig. S8a), 10 µM (Fig. S8b) or 20 µM (Fig. S8c) R5F2.

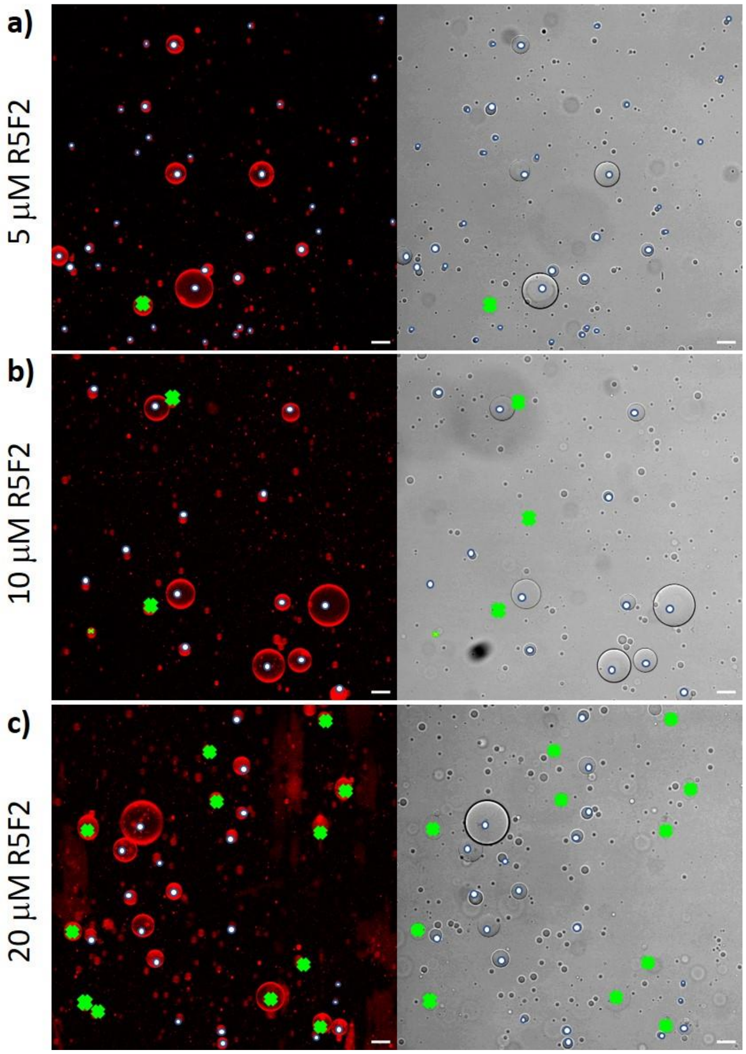


Fig. S8. Images of PRO' GUVs recorded on a Leica SP8 confocal microscope. In red (left): confocal cross sections; in grey (right): phase contrast, displaying enhanced contrast caused by sugar asymmetry across the membrane. The images were enhanced to more easily display the variations of phase contrast. Scale bar 50 µm. The GUVs marked with a white circle in the fluorescence channel show that they are also visible in the phase contrast image, meaning that they have not become leaky more than 10 minutes after the addition of the peptide. The crosses show GUVs that are visible in the fluorescent channel but not the bright field channel. a) PRO' + 5 µM R5F2; b): PRO' + 10 µM R5F2; c) PRO' + 20 µM R5F2.

The microscopy results reveal that the GUVs maintain their structural integrity and do not form flat contacts, indicating that the peptides do not induce adhesion. Additionally, the findings (Figs. S6 and S7) show that the membrane exhibits overall stability under the conditions tested, as evidenced by the finding that the GUVs remain intact.

Furthermore, no significant morphological changes are observed (Figs. S6-S8), apart from slight deflation due to sample evaporation or osmolarity mismatch, suggesting that peptides insert into the bilayer symmetrically in both leaflets without inducing substantial spontaneous curvature or measurable membrane area changes.
